# An intersectional expression platform for gene complementation using RNA-fragment end joining (REJ)

**DOI:** 10.64898/2026.09.13.751279

**Authors:** Lukas C. Bachmann, Ryan H. Hsu, Kip J. Hermann, Claire E. Williams, Richard L. Farman, Nicolas Criales, Craig Clark, Stella Kramer, Carolina Thörn Perez, Sawyer Randles, Karen Lettieri, Samuel L. Pfaff

## Abstract

Gene complementation is a powerful tool for genetic selection and protein functional studies but typically requires extensive screening for complementary components. Here we engineered a platform for gene complementation based on RNA end-joining (REJ) that precisely and efficiently splices separate RNA units into functional coding mRNAs. REJ is mediated by short structured modular RNA segments of ∼250bp that: (a) promote RNA::RNA interaction, (b) coopt the intrinsic cell splicing machinery to facilitate RNA trans-splicing, (c) minimizes translation of protein fragments from un-spliced RNA segments, and (d) encodes scar-free protein. We demonstrate that REJ is a broadly applicable system that can reliably split nearly any gene into complementary segments regardless of protein structure. To enable the use of this system we provide a toolbox of reporters and a web-based design tool to facilitate vector design. Demonstrated REJ applications include genetic complementation, intersectional labeling of cell types, and efficient expression of large proteins.

**Highlights:**

- Synthetic RNA cis-elements can direct efficient and precise RNA trans-splicing *in vivo*.
- Endogenous splicing factors catalyze RNA end-joining (REJ) without foreign proteins.
- The REJ platform can be applied to nearly any gene with few positional constraints.
- REJ can be used as an intersectional tool to build complex genetic logic circuits.

## Introduction

Many mechanistic studies and clinical diseases have a genetic component, which has led to the development of innovative approaches to modulate gene expression, repair mutations, and/or replace proteins ^1–7^. These successes have demonstrated that targeted control of gene expression and protein function have enormous significance for both research and clinical applications. Here we describe a system based on RNA::RNA reactions that can be used to create gene expression “logic circuits” that can be used for targeting cell types while minimizing the risk of immune system activation and off-target interactions.

RNA splicing is mediated by the ribonucleoprotein spliceosome complex present across cell types in higher eukaryotes^8^. The spliceosome catalyzes a trans-esterification reaction that joins exons within the same linear transcript and removes intervening intronic sequences via cis-RNA splicing (Figure 1A). RNA splicing is a well-established control point for setting gene expression levels and generating protein isoforms^9,10^. In contrast, the splicing together of separate non-linear RNAs is avoided in mammalian cells, possibly to minimize the risk of generating scrambled mRNAs that encode nonsensical proteins. Although trans-RNA splicing is not readily found in most species, organisms such as *C. elegans* do splice together separate RNAs in order to append leader sequences to their transcripts^11^. This is achieved by bringing the individual RNAs together using small nuclear ribonucleoproteins, raising the possibility that analogous trans-RNA esterification reactions may be possible to synthetically engineer in mammalian cells^12–14^. Indeed, recent studies have demonstrated that separate RNAs can be trans-spliced using endogenous splicing factors within mammalian cells if the two transcripts contain canonical splicing cis-elements and are brought together via Watson-Crick base pairing to form a pseudo-linear RNA substrate (Figure 1A) ^14,15^. These findings indicate that the mammalian cellular components for RNA splicing do not have a strict processivity requirement for translocation along a single continuous RNA strand.

**Figure 1:**
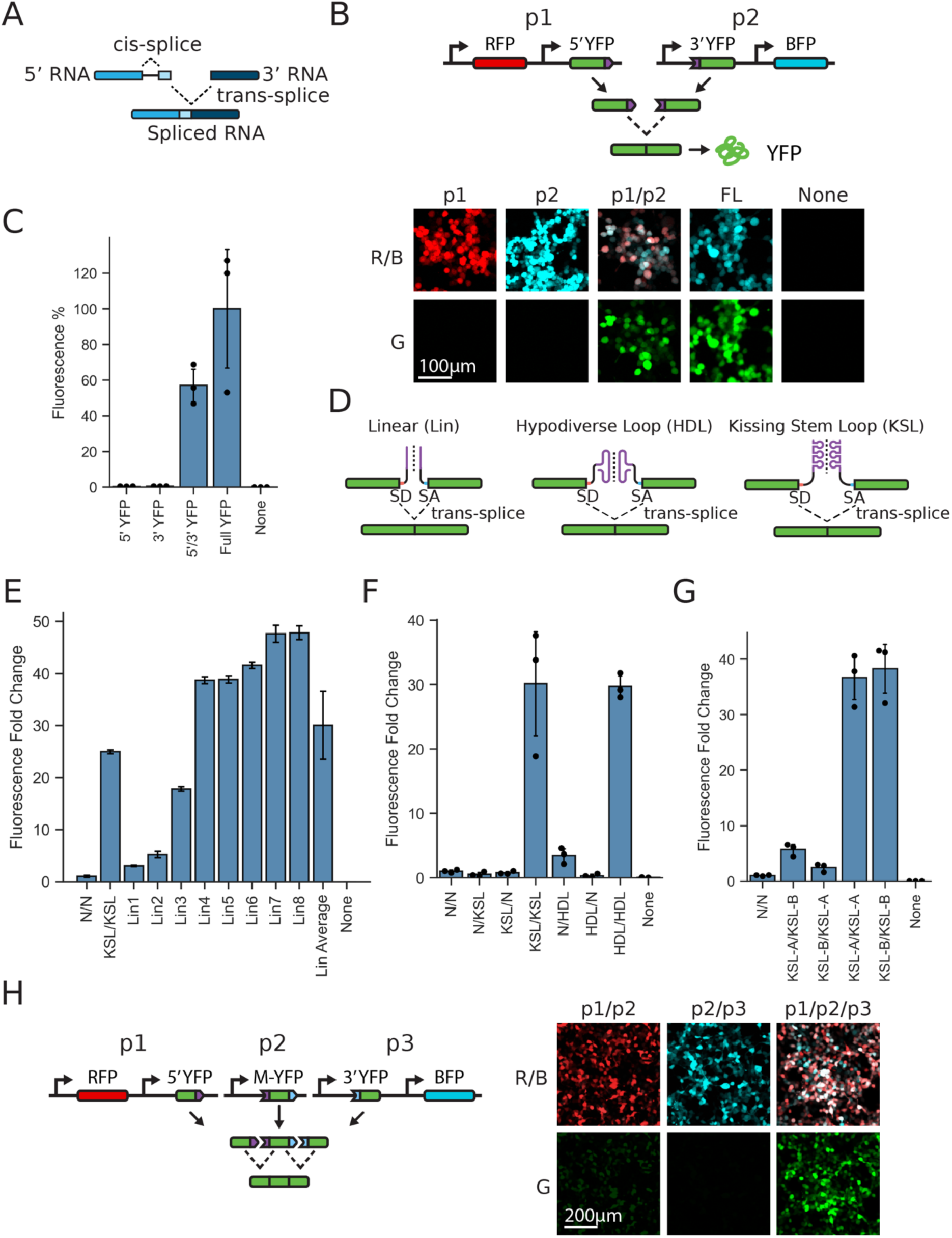
Structured RNA dimerization domains for trans-splicing. **A)** Schematic of RNA intramolecular cis-splicing versus intermolecular RNA trans-splicing. **B)** Fluorescence microscopy in HEK293T cells. YFP is split into two segments encoded on separate plasmids (p1 and p2) with RFP and BFP respectively to monitor cell transfection. Full-length YFP- and BFP-expressing plasmids served as positive expression controls. Neither the 5’YFP nor 3’YFP segments alone encode functional fluorescent protein, whereas co-transfection of p1 and p2 leads to YFP expression comparable to a plasmid encoding full-length YFP. Yellow fluorescent protein (YFP), blue fluorescent protein (BFP), red fluorescent protein (RFP). **C)** Quantification of YFP fluorescence following transfection of constructs shown in (B). YFP levels from co-transfection of p1 and p2 were 57.04 +/-11.11% compared to a full length YFP construct (p=0.0124). **D)** Schematic of RNA dimerization domains with Kissing Stem Loop (KSL), Hypodiverse Loop (HDL), and Linear (Lin) architectures for RNA::RNA interactions that facilitate trans-splicing. Canonical splice donors (SD) and splice acceptors (SA) were included to define exon-intron boundaries. **E)** Screen of multiple split-YFP reporters with different linear dimerization domains (Lin-RDDs). A 3-47 fold range of YFP reporter expression was detected across Lin 1-8. Split YFP reporters with KSL-RDDs or no binding domain (N/N) were included for comparison. **F)** Split-YFP reporter expression across RDD combinations. Matched dimerization structures mediate RNA trans-splicing, whereas mismatched or non-pairing configurations remain near background. Thus, complementary RDDs are selective and represent a key requirement for efficient trans-splicing (n=2-3 across groups; HDL/HDL vs N/N, p=0.0013; KSL/KSL vs N/N, p=0.0365). **G)** Split-YFP reporter expression across different combinations of KSL-RDDs. Matched pairs of complementary RDDs (KSL-A/A and KSL-B/B) expressed YFP, whereas mismatched combinations (KSL-A/B and KSL-B/A) were inactive (n=3 per condition; matched-vs-mismatched contrasts 6.4- to 15.6-fold; p=0.0054-0.0070). **H)** Trans-splicing of three separate RNAs. Schematic of plasmids with YFP divided into three segments termed 5′-YFP, Middle (M-YFP), and 3′-YFP. YFP fluorescence was detected when all three segments of YFP were co-expressed following HEK293T transfection.

RNA trans-splicing has many applications in biology. For example, viral vectors such as adeno-associated virus (AAV) have limited cargo capacity but are capable of co-infecting cells with high multiplicity. Thus, encoding sub-segments of a single gene in different AAV vectors that coinfect cells and transcribe separate RNAs that are spliced together to generate a single long mRNA is promising for expressing large proteins with vectors that have limited capacity^16^. Furthermore, RNA trans-splicing can be used to split genes into multiple non-functional components for gene complementation using intersectional genetics. This strategy can be used to create synthetic gene circuits and to target very specific cell populations while reducing the number of reporters needed. Although RNA trans-splicing has many potential applications, it has not been well characterized to establish a reliable platform system for piecing together fragments of any gene efficiently with minimal off-target activity.

To create an efficient and predictable combinatorial system for gene complementation as a tool for intersectional genetics we created short RNA motifs of ∼250 nucleotides that can be appended to transcripts to direct RNA trans-splicing. These motifs mediate RNA end-joining (REJ) of separate RNAs by recruiting the intrinsic cellular splicing machinery for the trans-esterification reaction, which we tested and optimized to be highly efficient while maintaining the nucleotide joining precision and minimizing off-target interactions. Because this system is based on RNA and uses only intrinsic cellular proteins it avoids the introduction of potential antigens. We designed RNA::RNA interaction modules containing splice enhancers that promote efficient RNA joining. We demonstrate that REJ can express large therapeutic genes, perform intersectional genetics, and implement complex logic circuits *in vivo*. Together, our findings establish REJ as a widely applicable platform technology that functions broadly across genes and cell types.

### Design

Mammalian cells are capable of splicing together separate RNAs if the two strands base pair and contain cis-elements for splicing ^15,17–19^. However, a reliable vector system that is highly efficient, has documented low off-target activity, suppresses the translation of protein fragments, functions across different cell types, and can be programmed with orthogonal components to promote the trans-splicing of more than two transcripts has not been established. To overcome this limitation, we developed a modular platform called RNA end-joining (REJ), which has broad applications in gene complementation, intersectional genetics, and the expression of large proteins. REJ modules ≤250 nucleotides in length were screened for their ability to direct efficient and precise intermolecular RNA joining across cell types and transgenic constructs. The REJ modules were optimized to perform four coordinated functions: promote selective RNA interactions through structured complementary sequences; recruit the endogenous spliceosome using canonical splice donor and acceptor elements to catalyze trans-splicing; suppress translation from unspliced RNA intermediates to minimize the accumulation of truncated protein products; and generate scar-free, mature mRNAs that encode native, full-length proteins without residual amino acid sequences at the junction. Because REJ relies exclusively on endogenous splicing machinery, the platform eliminates the need for exogenous recombinases, inteins, ribozymes, or other foreign proteins. Its modular architecture enables implementation of intersectional genetic logic by restricting functional gene expression to cells that co-express all required transcripts. These features enable the design of increasingly complex genetic circuits for temporal and spatial control of gene expression. To facilitate the use of this platform, we provide REJ vectors, a toolbox of split reporters, a design algorithm and associated web tool REJ Studio, to aid in the design of novel vectors.

## Results

### Characterizing elements for RNA::RNA interactions for trans-splicing

To examine the feasibility of creating a reliable system for combinatorial genetics using trans-splicing of RNAs, we split YFP into two non-functional segments that are transcribed from separate plasmids (Figure 1B). The N-terminal YFP coding 5’-RNA incorporated synthetic sequences of 219 nucleotides at the 3’ end containing multiple elements: consensus splice donor (SD) and intronic splice enhancer (ISE) elements identified from alignments of intron-exons^20^, an RNA dimerization domain (RDD), a polyadenylation signal sequence, and a separate RFP reporter to detect cell transfection (Figure 1B). The C-terminal YFP 3’-RNA had a 5’ 232 nucleotide sequence containing: a complementary RDD, intronic splice enhancer elements, a poly-pyrimidine tract for RNA-branch formation, a canonical splice acceptor (SA), and a separate BFP reporter to detect cell transfection (Figure 1B). Separately, neither 5’ nor 3’ REJ YFP RNAs were capable of encoding fluorescent protein; whereas the combination of 5’ and 3’ REJ YFP RNA constructs expressed significant levels of YFP, albeit slightly less than from a single full-length construct (Figure 1B-C).

We and others have tested different complementary sequences for the RDD elements and observed a wide range of trans-splicing efficiency^18,21–23^. We reasoned that the secondary structure of the RDD may attenuate RNA::RNA interactions and limit the efficiency of trans-splicing. RNA secondary structure can be difficult to predict in long RNAs and would presumably change depending on the particular transgene being expressed. Therefore we tested multiple RDD sequence designs in order identify motifs that facilitate efficient trans-splicing across a range of transgenes.

We screened a set of random complementary “unstructured” linear (Lin) 50-mer RDD sequences and observed a range of trans-splicing efficiencies (Figure 1D, E). Because the variability in efficiency of Lin-RDDs might be due to the influence of flanking sequences, it was unclear if a single specific Lin-RDD would function in the context of different transgenes. This prompted us to test whether structured RDDs were functional alternatives to Lin-RDDs. First we created structured RDDs with a hypodiverse loop (HDL) for base pairing (Figure 1D). Namely, the hypodiverse RDD regions were 100bp segments with only pyrimidines or the complementary purine nucleotides. This design was predicted to reduce intra-molecular interactions that could occlude dimerization of the RNAs. RDD motifs with the HDL design expressed YFP at levels comparable to the high-performing Lin constructs (Figure 1E, F).

Next we examined whether highly structured RDDs can facilitate RNA trans-splicing. We designed RDDs with stem loop sequences based on the kissing stem loops (KSL) identified within HIV for RNA dimerization ^24^ (Figure 1D). We found that RNAs with complementary KSL structures were efficiently trans-spliced (Figure 1E-G). Moreover, the RDD segments of the KSLs could be “programmed” to have specificity for splicing to the appropriate complementary RNA (Figure 1G). Using orthogonal KSL-RDDs we found that three RNAs could be joined in the proper order and orientation (Figure 1H). These findings indicate that it may be possible to assemble very long mRNAs encoding large proteins from ≥2 RNAs. In addition, orthogonal RDDs can be used to create complex multi-layer intersectional logic circuits for controlling gene expression.

Taken together, our findings reveal that a variety of RDDs can facilitate trans-splicing of RNAs. We have not systematically compared each RDD across many genes, but each design is capable of expressing similar levels of YFP. We note, however, that we have used the KSL-RDD to efficiently express >25 different split genes (see below), and therefore the structured conformation of this RDD may minimize negative interactions with flanking sequences. Thus, we found that without further optimization the KSL-RDD design yields reliable activity in many different contexts.

### Cis-elements for efficient trans-splicing

The vectors we designed to mediate RNA trans-splicing were based on inclusion of multiple canonical cis-elements recognized by the cell’s intrinsic spliceosome ribonucleoprotein machinery (Figure 2A). To establish the functional contribution of each cis-element for trans-splicing we quantified gene expression from vectors with specific mutations. As expected, mutation of the canonical SD and SA sequences blocked YFP expression (Figure 2B). Both the 5’ and 3’ RNAs in the split YFP vectors included canonical intronic splicing enhancers (ISEs) thought to actively increase splicing efficiency across many cell types^25–27^. Mutation of the splice enhancer elements in the 5’ RNA had a greater impact on YFP expression than the 3’ RNA splice enhancers in HEK293T cells (Figure 2C).

**Figure 2:**
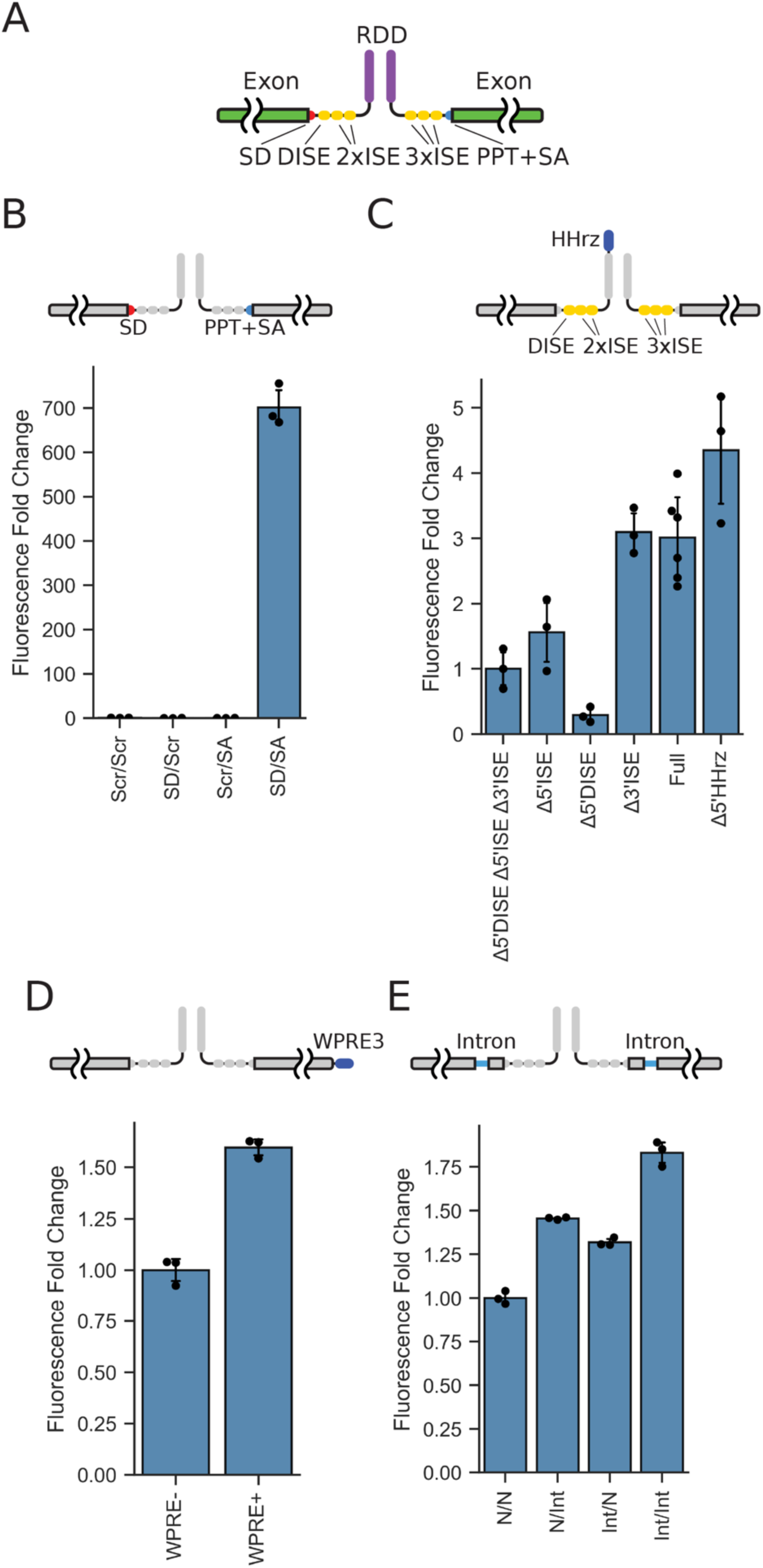
Optimization of cis-elements for expression of trans-spliced mRNA. **A)** Schematic of modular REJ components and their relative topology within two RNAs encoding a split genetic cargo. Splice donor (SD), downstream intronic splice enhancer (DISE), array of unique intronic splice enhancers (2x or 3xISE), poly-pyrimidine tract (PPT), splice acceptor (SA). **B)** Split-YFP reporter expression was quantified after disruption of canonical splice donor and splice acceptor motifs. Intact SD/SA motifs drove robust YFP expression, whereas mutation (replacement with scrambled sequence, Scr) of either splice site abolished reporter expression. n=3 per condition; intact SD/SA vs SD/Scr and Scr/SA, p=0.0015 for both; up to 2283-fold difference. **C)** Loss-of-function deletions in cis-elements to quantify their contribution toward mRNA trans-splicing and reporter expression. Deletion of the downstream intronic splice enhancer (Δ5’DISE) reduced reporter expression 10.4-fold compared to the intact system, p=0.0001. Deletion of the 5’ intronic splice enhancer (Δ5’ISE) lowered expression 1.9-fold, p=0.0182. Deletion of all splice enhancers (Δ5’DISE, Δ5’ISE, Δ3’ISE) reduced expression 3.0-fold, p=0.0005. Designs containing all elements (Full) or Hammerhead ribozyme (HHrz) shown. Thus, splice enhancers increase reporter expression from trans-spliced mRNAs (n=3-6 across groups). **D)** The addition of woodchuck hepatitis virus posttranscriptional regulatory element (WPRE) on the 3’ RNA increased YFP reporter expression 1.6-fold relative to the control lacking a WPRE (n=3 per group; p<0.001). **E)** Split-YFP constructs containing short cis-introns in one or both 5’/3’ RNA segments stimulated reporter expression: no cis-intron (N), inclusion of cis-intron (Int). Relative to the control N/N, reporter expression increased 1.32-fold for Int/N (p<0.001), 1.46-fold for N/Int (p=0.0016), and 1.83-fold for Int/Int (p<0.001) (n=3 per condition).

The woodchuck hepatitis virus post-transcriptional regulatory element (WPRE) increases transgene expression from a variety of vectors by facilitating polyadenylation (transcript termination), increasing nuclear export of mRNA, and stabilizing transcripts^28^. The addition of a WPRE element in the 3’ RNA increased YFP expression by ∼1.5 fold (Figure 2D). Furthermore, RNA splicing stimulates gene expression at multiple cellular levels and therefore transgenic vectors often include introns if space permits^25,29,30^. We found that the addition of cis-introns within both the 5’ and 3’ RNA segments markedly enhanced the overall level of transgene expression from the trans-spliced vectors (Figure 2E).

In addition to incorporating elements for increasing the overall level of gene expression, we included sequences that attenuate the expression of protein fragments from un-spliced RNA segments. While some applications with trans-splicing may not be impacted by the expression of truncated proteins, we aimed to minimize the expression of truncated proteins from the un-spliced RNAs to avoid expressing peptides with dominant negative activity. To reduce the translation of un-spliced 5’ RNA we included a hammerhead ribozyme element (HHrz) to target degradation of the un-spliced 5’ transcript (Figure 2C). The 3’ RNA included out-of-frame ATG codons as decoy translation start sites to limit initiation from internal ATGs within the transgene protein.

Taken together, we defined a set of cis-elements that facilitate gene expression at multiple post-transcriptional levels from trans-spliced mRNAs. These features include elements for RNA dimerization, splicing, polyadenylation, RNA stability/transport, and translation. We termed this process RNA end-joining (REJ). Below we demonstrate that the compact synthetic group of cis-elements that mediate REJ can be used to split nearly any mRNA and is functional in many tissues *in vivo*. Since REJ modules coopt the cell’s intrinsic splice machinery, no foreign proteins are required, thereby reducing the risk of triggering an antigenic response. Importantly, following trans-splicing all REJ elements are removed, thereby generating an mRNA that encodes only the transgenic protein of interest without internal “scars” or additional peptide sequences.

### REJ efficiency

The efficiency of transgenic vectors is important for achieving protein levels that are functional with lower titers of viruses. We compared the level of REJ-mediated YFP expression encoded by coinfection with dual AAV vectors *in vivo* to the level of YFP from a single vector using the same promoters, capsids and ORF-matched concentrations. Muscle cells injected with dual REJ AAV vectors expressed >80% of the YFP fluorescent levels observed with a single vector (Figure 3A). Several DNA-, RNA-, and protein-based approaches have been developed to overcome the cargo size limits of AAV based on splitting genes into multiple virus particles^17,31,32^. In comparing the efficiency of split vectors using either REJ or DNA-recombination to express YFP in mouse muscle, we observed approximately 15-fold higher YFP fluorescence with dual REJ vectors compared to dual DNA-recombination vectors^17^ (Figure 3B).

**Figure 3:**
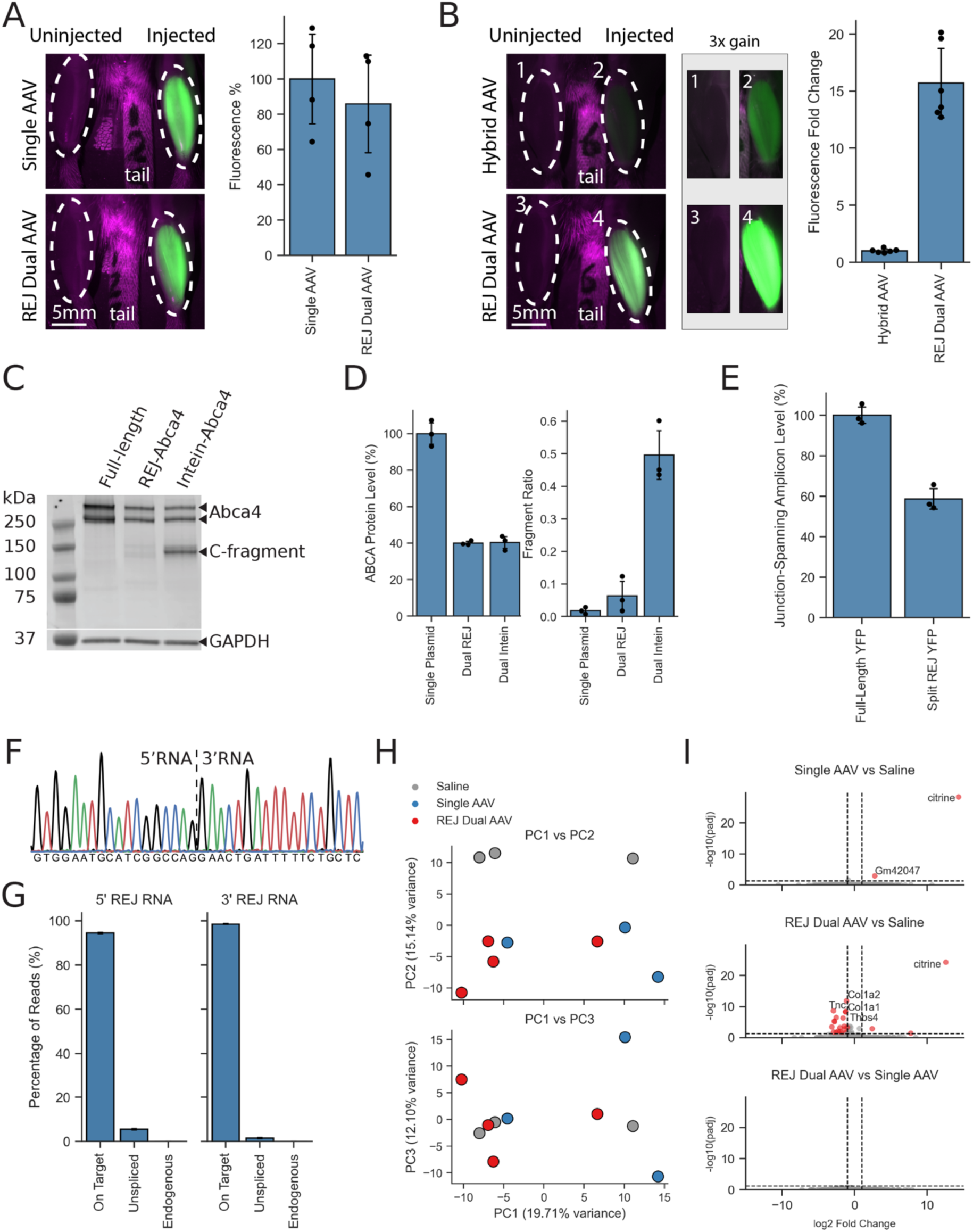
Analysis of REJ-mediated mRNA trans-splicing efficiency, precision, and off-target interactions. **A)** *In vivo* YFP fluorescence was measured in mouse tibialis anterior (TA) muscle after injection of saline, single AAV, or dual AAVs expressing split YFP mRNAs with REJ modules. Four weeks following unilateral viral injection REJ dual AAVs expressed YFP at ∼80% the level of the control single AAV (n=2-4 across groups). Uninjected TA muscle also shown. **B)** Comparison of YFP fluorescence in mouse TA muscle four weeks following unilateral limb injection of AAVs. REJ dual AAV expressed 15.7-fold higher YFP fluorescence than split-YFP Hybrid AAV vectors (DNA-recombination). 3x gain increase shows YFP expressed by Hybrid AAV vectors and the saturated signal from REJ dual vectors. n=6 per group; p<0.001. **C)** Western blot of 250-256 kDa ABCA4 protein expressed using REJ and intein vector systems in transfected HEK293T cells. Full-length corresponds to control expression from a single plasmid encoding ABCA4. Full length ABCA4 protein (appearing as a doublet) is expressed by both REJ and intein vectors systems. The anti-C-terminal ABCA4 antibody detects a prominent C-terminal fragment of ABCA4 encoded by the intein vectors as expected, but not the REJ vectors. GAPDH is shown as a loading control. **D)** Full-length ABCA4 protein levels were quantified for HEK293T cell transfections with Single Plasmid, Dual REJ, and Dual Intein constructs. REJ- and intein-vectors expressed full-length ABCA4 similarly at ∼ 40% of Single Plasmid controls (n=3 per group; Dual REJ vs Dual Intein p=0.8988). Dual REJ vectors expressed a 7.8-fold lower fragment-to-full-length protein ratio than Dual Intein vectors (n=3 per group; Dual REJ vs Dual Intein p=0.0045). **E)** qPCR quantification of YFP trans-spliced mRNA levels expressed from dual split-YFP REJ vectors relative to levels expressed by a single full length YFP vector control. REJ-vectors expressed 58.7% the level of spliced RNA compared to the control (n=3 per group; p=0.0010). **F)** A Sanger sequencing chromatogram spanning the REJ junction shows RNA trans-splicing occurs precisely at the CCAG|GAAC splice junction (SD/SA junction) as expected, thereby ensuring all REJ elements are removed following trans-splicing, generating a “scar free” ABCA4 mRNA. **G)** Following infection of mouse tibialis anterior muscle with dual REJ-YFP AAV vectors, RNA was purified from pooled samples and sequenced. Transcripts were classified as on-target (joining of the 5’ and 3’ YFP RNAs at the correct SD/SA junction), unspliced, or off-target splicing of either the 5’- or 3’-YFP RNAs to an endogenous cellular transcript. Most classified pooled reads were on-target (5′-YFP: 4954/5242, 94.51%; 3′-YFP: 4943/5018, 98.51%), and the remaining classified reads were unspliced (5′-YFP: 288/5242, 5.49%; 3′-YFP: 75/5018, 1.49%). Off-target splicing of either 5′- or 3’-YFP RNAs to cellular transcripts was not detected (0 observed in both datasets; 95% upper bound <0.0572% for 5′-YFP and <0.0598% for 3′-YFP). **H)** Principle component analysis (PCA) of transcriptomes from TA muscle injected with Saline, Single-YFP AAV, and REJ Dual-YFP AAVs. PC1 correlated with batch, while PC2 correlated with viral versus non-viral infected samples. Transcriptomes from Single and REJ Dual AAV samples were intermixed in PCA space (n=3-4 across groups; PC1 19.71%, PC2 15.14%, PC3 12.10% variance explained). **I)** Volcano plots of all pairwise muscle-transcriptome comparisons among Saline, Single-YFP AAV, and REJ Dual-YFP AAV injected samples found few differences between single and dual vector systems. As expected, citrine expression was robustly detected in the AAV infected mice, whereas a REJ-specific transcriptomic alteration was not detected. Significance thresholds are padj<0.05 and |log2 fold change|>1 (20,616 genes tested per comparison).

Next we compared the expression levels of the large ABCA4 protein using REJ or inteins to reconstitute the protein from split vectors. *ABCA4* encodes a 256 kDa retinal membrane protein mutated in Stargardt disease whose mRNA exceeds the capacity of a single AAV^33^. Screening has led to the identification of intein-based dual AAV ABCA4 vectors that express efficacious levels of protein for treating Stargardt disease^34,35^. We detected similar levels of full length ABCA4 protein expression with both REJ and intein vectors (Figure 3C, D). However, because inteins function by catalyzing the ligation of protein fragments, we found the ratio of fragmented protein to full-length ABCA4 was approximately 8-fold reduced with REJ (Figure 3C, D).

In addition to intein-based approaches, several dual-vector strategies have been developed to overcome the packaging limitations of AAV, including ribozyme-mediated mRNA trans-ligation (STITCHR) and Cre recombinase–dependent DNA reconstitution (AAVLINK) ^36,37^. A key question is how the efficiency of REJ compares with other split-gene technologies that generate full-length mRNAs. Direct comparisons of protein output across dual-vector systems are challenging because expression levels depend on numerous factors, including viral titers, vector stoichiometry, and the conformation and subcellular localization of the split components. These variables are difficult to standardize across different constructs and transgenes. To enable a more comparable assessment across platforms, we quantified full-length mRNA production from dual REJ vectors relative to a single-vector construct expressing the same reporter. This metric provides a normalized measure that can be compared across genes, cell types, and split-gene technologies.

Using this approach, we found that REJ consistently generated approximately 50% of the full-length mRNA produced by single-vector controls across a diverse range of cell types and transgenes (Figure 3E; see below). Measurements of REJ-mediated RNA trans-splicing efficiency closely matched protein expression levels, with dual REJ vectors producing ∼50% of the protein output observed with single-vector controls (Figure 4C, D). These results suggest that both existing^17,31,32,36,37^ and future split-gene technologies are unlikely to exceed REJ efficiency by more than approximately two-fold, as a single-vector system represents the theoretical upper limit for expression.

**Figure 4:**
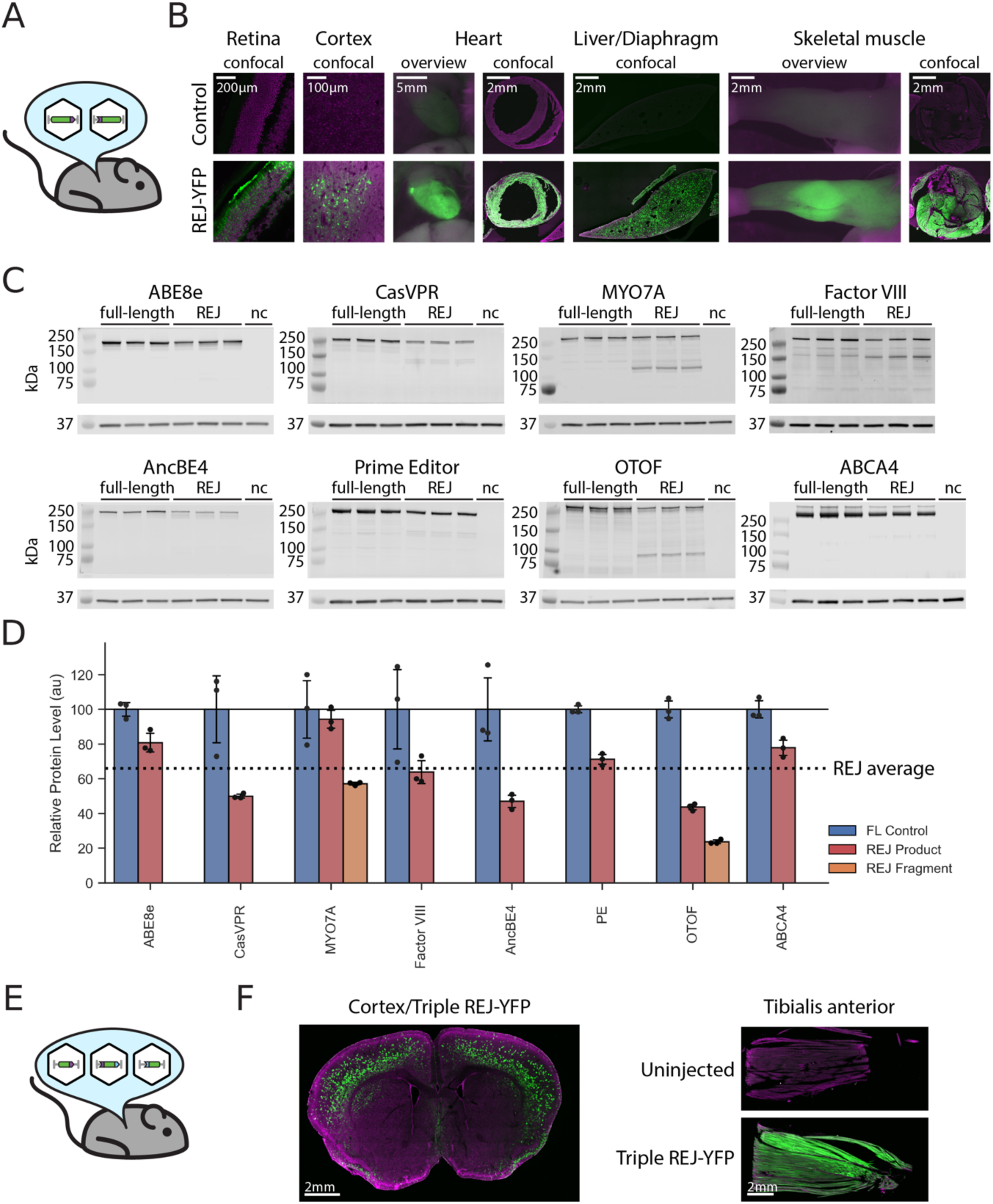
REJ is a platform for gene expression. **A)** Schematic illustrating *in vivo* delivery of split-YFP dual vectors with REJ motifs. **B)** YFP fluorescence imaging with a combination of whole tissue (overview) and confocal microscopy of tissue sections. Controls are saline injected samples. (Left to right) REJ vector expression is detected in: retinal photoreceptors, cortical cells, heart tissue, liver and diaphragm muscle in same section, and tibialis anterior skeletal muscle. Scale bar size noted on each image. **C)** Western blots of HEK293T transfected with a full-length positive control or two plasmids containing split genes with REJ motifs for: Abe8e, Cas9-VPR, Myo7a, Factor VIII, AncBE4, Prime Editor, Otoferlin, and ABCA4. Several of the REJ constructs (i.e. Myo7a) lacked the cis-elements that suppress translation of unspliced RNA (5’ RNA hammerhead ribozyme element and 3’ RNA out-of-frame ATGs), whereas some REJ constructs included the cis-elements that suppress translation of protein fragments (i.e. ABCA4). In each case full length protein was expressed by the split REJ vectors. Non transfection control (nc). **D)** Full length (FL), REJ product, and REJ protein fragment levels quantified from Western blot in (C) (prime editor, PE). Across all tested examples the dual REJ system on average expressed 66.06% +/-6.42% the level of protein compared to a single (full length) control. **E)** Schematic of YFP split into three AAV vectors with REJ motifs containing orthogonal binding domains designed to pair the RNAs in the correct order and orientation to reconstitute full length reporter expression *in vivo* via multiple trans-splicing steps. **F)** Confocal imaging detects YFP fluorescence in the mouse cortex and tibialis anterior muscle four weeks after injection of the three AAVs (schematic in E) into these tissues. YFP shown in green, contrast autofluorescence shown in magenta.

### REJ precision and off-target interactions

To ensure that RNA splicing mediated by REJ occurred at the proper SD/SA junctions, we sequenced RNA from transfected HEK293T cells. As expected, RNAs were joined at the correct site indicating splicing-fidelity was not significantly altered by joining separate RNAs (Figure 3F).

To determine if REJ modules are prone to off-target splicing with endogenous cellular RNAs we infected mouse muscle with AAVs expressing the 5’-REJ and 3’-REJ RNAs encoding YFP and sequenced the cellular transcripts. Most REJ RNAs were spliced together, while a small proportion of REJ RNAs were un-spliced (Figure 3G). Importantly, we did not detect YFP transcripts spliced to cellular RNAs (Figure 3G, Methods).

To further test the system with an alternative transgene, we transfected HEK293T cells with high amounts of plasmid encoding only a single 5′ or 3′ REJ-Abe8e RNA segment. This experiment was designed as an extreme scenario where the transcripts are expressed at atypically high levels without its complementary REJ-RNA splicing partner. We then performed deep sequencing yielding 83 million (5’ REJ-RNA) and 78 million reads (3’ REJ-RNA). Candidate splice products were required to have at least five supporting reads to reduce singleton and sequencing-error calls. After filtering, we identified 5,394 reads in the 5′-only condition and 38,239 reads in the 3′-only condition that contained the REJ anchor sequence. None of the 5′ REJ-Abe8e RNAs were detectably spliced to endogenous cellular transcripts. Rare 3′ REJ-Abe8e trans-splicing to endogenous transcripts was detected, with the most abundant endogenous splice product representing only 0.097% of classified 3′ REJ-Abe8e reads. These findings indicate that endogenous trans-splicing by REJ is exceedingly rare, even under highly exaggerated expression conditions, and is unlikely to deplete or perturb any specific cellular transcript.

Overt changes in the growth rate or appearance of cells transfected or infected with REJ vectors were not observed *in vivo* or *in vitro* (see Figures 1C; 3A). To detect cellular alterations more sensitively we examined whether REJ vectors encoding YFP triggered transcriptional changes. The transcript profile of muscle cells was compared across three conditions: uninfected (normal) muscle, muscle infected with a single AAV encoding YFP, muscle infected with dual

REJ-AAVs encoding YFP (Figure 3H, Methods). Principal component analysis of the transcriptomes revealed a difference in PC2 of uninfected tissue from AAV-infected muscle, which appears to have arisen as a reaction to the virus itself highlighting the sensitivity of this assay. Because the principal components of the single AAV and dual REJ-AAV samples were intermixed, the transcriptome of cells did not appear to be markedly altered by REJ-mediated trans-splicing.

Next we examined whether the double strand (ds) RNA formed by the RDD segments in the REJ modules activate pattern recognition receptors (PRRs) such as TLR3, RIG-I and PKR that sense cytoplasmic dsRNA^38,39^. While PRRs are expressed in muscle tissue^40^, we failed to detect the expression of downstream transcripts associated with activation of the dsRNA sensors in muscle infected with REJ vectors (Figure 3I). Although it is unclear how REJ evades the PRR system, perhaps the short length of the complementary RDD segments within structured kissing domains and the nuclear localization of the paired REJ RNAs are contributing factors. Taken together, the REJ system did not trigger overt cytotoxicity based on transcriptomic, morphological, or growth patterns of cells.

### REJ is active across cell types and genes

We examined whether the REJ system is active within different cell types using AAV8 REJ-YFP vectors with the ubiquitous CMV promoter. We found that systemic delivery of the dual AAV8 REJ vectors in adult mice led to YFP expression in many tissues including the diaphragm, cardiac muscle, skeletal leg muscles, and liver (Figure 4A, B). Likewise, subretinal injection labeled retinal neurons whereas injection into the cortex labeled cortical neurons and surrounding cells (Figure 4B). In summary, we observed YFP in every mouse tissue type tested when the REJ system was combined with the appropriate promoter, capsid and injection method. These findings are consistent with the likelihood that all cell types with the machinery for mRNA splicing are capable of RNA trans-splicing mediated by REJ modules.

To determine if the REJ vector system could be useful as a reliable platform technology to reconstitute the expression of different genes without special optimization we created dual plasmid vectors for 8 large proteins that exceed the coding capacity of AAV (Figure 4C). Across all genes, transfected cells expressed 66.1 ± 6.4% the level of protein from dual REJ vectors compared to controls (4C, D). The subset of REJ vectors that incorporated sequences to reduce N- and C-terminal protein fragment translation (see above) expressed predominantly full-length protein (e.g. ABCA4; Figure 4C, D). Thus, the REJ system is a relatively efficient expression system for many different genes without the need for large scale screening to optimize.

In some instances full length cellular proteins exceed the capacity of two AAV vectors. Because the RDD region of the REJ motif can be programmed to have a specific complementary partner we identified orthogonal RDD pairs (see Figure 1G) that can be used in combinatorial patterns. YFP was split into three segments and REJ motifs with two different RDD types were used to bridge the three RNAs in the proper order and orientation. AAVs encoding the three segments of RNA encoding YFP were injected into the mouse cortex and tibialis anterior muscle (Figure 4E). Without amplification YFP expression was detected in both cortical and muscle tissue (Figure 4F). These findings indicate that the REJ system can be used to join multiple RNA segments, however, the efficiency of protein expression will likely be reduced as the number of gene segments increases.

### Tools for intersectional-genetics

We generated a set of intersectional genetic tools based on REJ-mediated complementation for genes frequently used to study biological mechanisms (Table 1). This toolkit includes dual-vector systems for fluorescent and secreted reporters, including mCitrine/YFP, triple-part mCitrine/YFP, and Gaussia luciferase; functional reporters and actuators, including GCaMP, hM3Dq DREADD, and ReaChR; recombinases and regulatory proteins, including iCre, FlpO, TetOFF/tTA, and dCas9-VPR; selection and genome-engineering tools, including HSV-TK, ABE8e, AncBE4, and PE2 / Prime Editor; and disease-relevant large genes, including ABCA4, OTOF, MYO7A, and Factor VIII. We also generated REJ multiple-cloning-site backbone plasmids to facilitate custom split-gene designs. Expression or activity of REJ-reconstituted transgenes was validated in cell culture using fluorescence imaging, immunofluorescence, Western blotting, antibiotic selection, or functional reporter assays, and a subset of constructs was further tested in vivo. These plasmids are available through Addgene and were designed to facilitate cloning and AAV production.

REJ utilizes canonical splicing and therefore split sites must be selected at canonical WG|GW exon-exon boundary motifs where W represents a T or A nucleotide. Through synonymous codon substitution (Figure 5A), the candidate WG|GW split motifs are highly abundant across coding sequences (Figure 5B). Because recombination occurs at the level of RNA, split motifs can be selected regardless of their relation to codon-codon boundaries.

**Figure 5:**
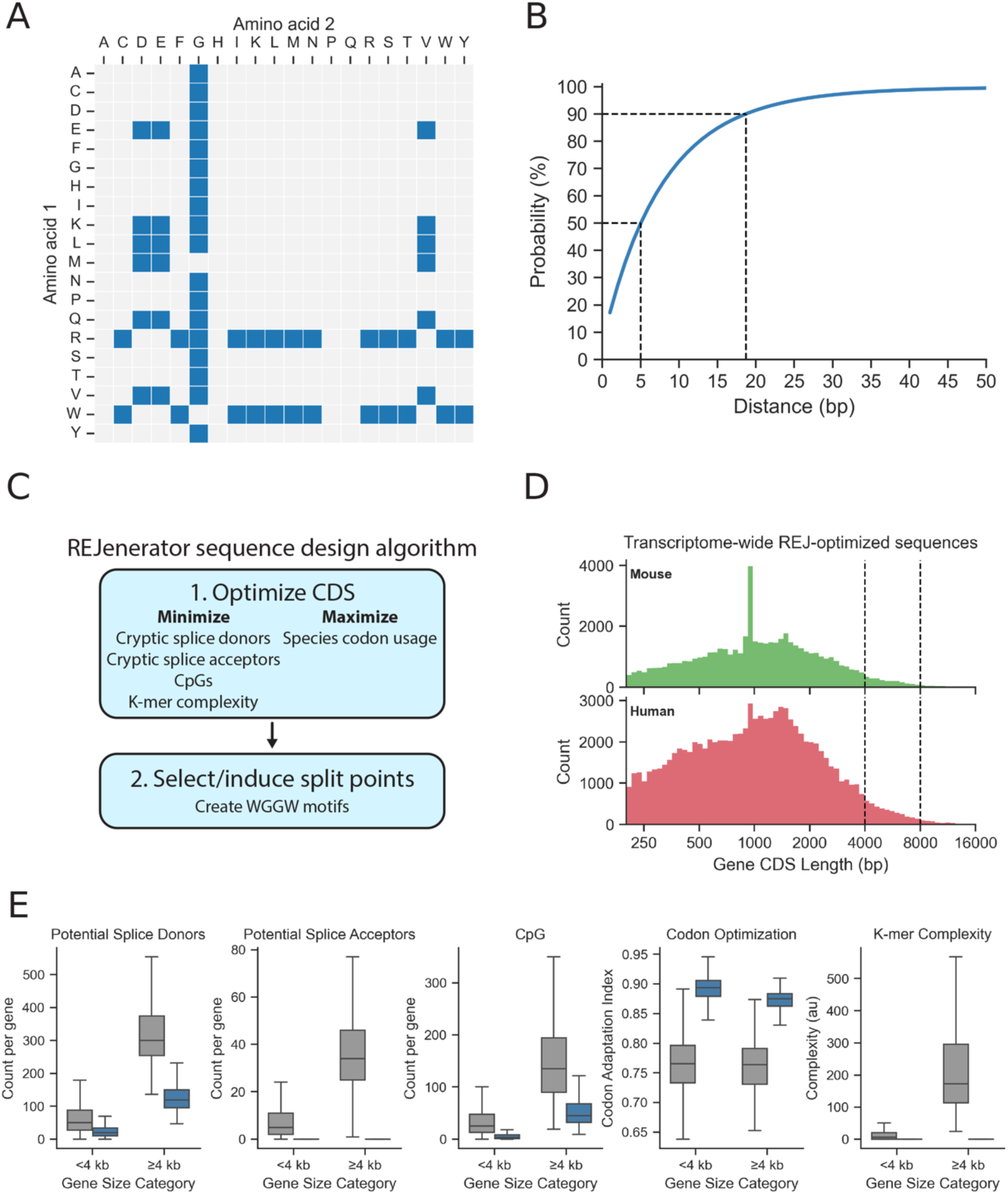
Computational tools for using REJ to split genes. **A)** REJ exon-exon boundaries are encoded by canonical WG|GW (W: A or T) motifs that are not constrained to codon-codon boundaries and therefore inducible through synonymous codon substitution. Across all amino acid pairs, 62 (blue) of 400 can be encoded using codons containing a WG|GW motif. **B)** A transcriptomic assessment of 10,000 human coding sequences was conducted to evaluate the feasibility of identifying REJ-compatible split points across diverse protein targets. This revealed high flexibility in splice-site selection during construct design, with a >50% probability of identifying an inducible WG|GW motif within 5 bp of any target coordinate and a >90% probability within 19 bp. **C)** Schematic of the REJenerator design algorithm. Starting from a naive coding sequence, REJenerator applies synonymous codon substitutions to reduce cryptic splice donor/acceptor motifs, tune GC content and k-mer complexity, and maximize species-specific codon adaptation (CAI). The optimized sequence is then scanned for split sites with a WG|GW motif for insertion of REJ modules and stimulatory introns, enabling rapid design of high-performance split REJ constructs. **D)** REJenerator was used to generate synthesis-ready REJ sequences of all 91,049 human and 52,809 mouse protein isoforms. Coding-sequence length distributions are shown. A total of 5,698 transcripts (3.96%) were ≥4,000 and <8,000 bp (requiring a 2-part REJ design), and 618 transcripts (0.43%) were ≥8,000 bp (requiring a 3-part REJ design). **E)** Boxplots of REJenerator constraint metrics before and after optimization, stratified by coding sequence length (<4,000 bp, n=137,542; ≥4,000 bp, n=6,314). Optimization reduced donor counts, acceptor counts, CpG counts, and k-mer complexity while increasing codon optimization (CAI) in both size classes.

Empirically, at any given position it is 50% likely to find a present or inducible (i.e. synonymous codon substitution) split site within 5 bp, and 90% likely to find one within 19 bp (Figure 5B). In our testing, we found that all WG|GW split sites have been compatible with robust REJ-mediated trans-splicing and subsequent protein expression.

To facilitate the design of optimized genetic systems that exploit RNA trans-splicing with REJ we created a two-stage sequence design algorithm (REJenerator) and an associated web application (REJ Studio). In the first stage, the algorithm simultaneously optimizes multiple objectives through synonymous codon substitution: (a) depletion of potential off-target splice donors and splice acceptors, (b) reduction of CpG dinucleotides, (c) control of k-mer complexity, and (d) species-specific codon optimization. These objectives are constrained by maintaining the desired peptide sequence and keeping GC content in the range of 35-60% (Figure 5C).

Therefore, the algorithm streamlines the process of minimizing potentially cryptic splicing, reducing immunogenicity, enhancing viral expression, facilitating protein translation, and simplifying DNA synthesis and cloning. In the second stage, the algorithm induces exon-exon boundary motifs in specific split regions to install the REJ modules and optional 5’ and 3’ stimulatory introns. These sequences are synthesis-ready and can be expressed through any standard expression vector. To facilitate research applications of REJ, we used REJenerator default parameters to design optimized REJ constructs across all annotated 91,049 human and 52,809 mouse protein isoforms, 4.4% of which exceed the cargo capacity of AAV (Figure 5D, E). We also compiled OMIM disease-associated genes with annotated coding sequences exceeding single-AAV cargo capacity to highlight large therapeutic targets that may benefit from REJ-based split-gene design (Supplemental Table 1). These optimized sequences, the disease-gene resource, and a web interface for designing custom sequences using the REJenerator algorithm with additional parameters are made publicly available on REJ Studio (www.rejstudio.com; during peer review the site is password-protected. See Supplemental Tutorial Video 1 and Figure S1).

### Gene complementation and combinatorial applications

It is advantageous to target specific cell types in the context of heterogenous cell populations for functional characterization. While numerous promoters/enhancers with cell type-selectivity have been identified, intersectional approaches are commonly used to achieve greater specificity using tools such as combinations of Cre and Flp^41^. The REJ system can potentially simplify intersectional experiments by expressing a single reporter in cells where two different promoters have overlapping activity. Cell targeting with REJ vectors can be further refined by selecting the appropriate capsid and viral delivery strategy.

We first split the transcription transactivator dCas9VPR into two segments and tested the intersectional activation of UAS transgene reporters with and without the appropriate guide RNA (Figure 6A). Controls with the UAS-reporter that lacked dCas9-VPR and the guide did not express detectable YFP, whereas cells with all three components expressed the reporter. Next we split the tetracycline activator (tTA) into two segments with REJ modules and show that coexpression of 5’-tTA and 3’-tTA strongly activates a Tetracycline-reporter (Figure 6B, Figure S2). The addition of doxycycline inhibits tTA and represses expression of the tetracycline-reporter. Together these findings reveal how dCas9VPR and tTA can be used to build intersectional gene control circuits that are gated by the presence of a guide RNA or absence of doxycycline, respectively.

**Figure 6:**
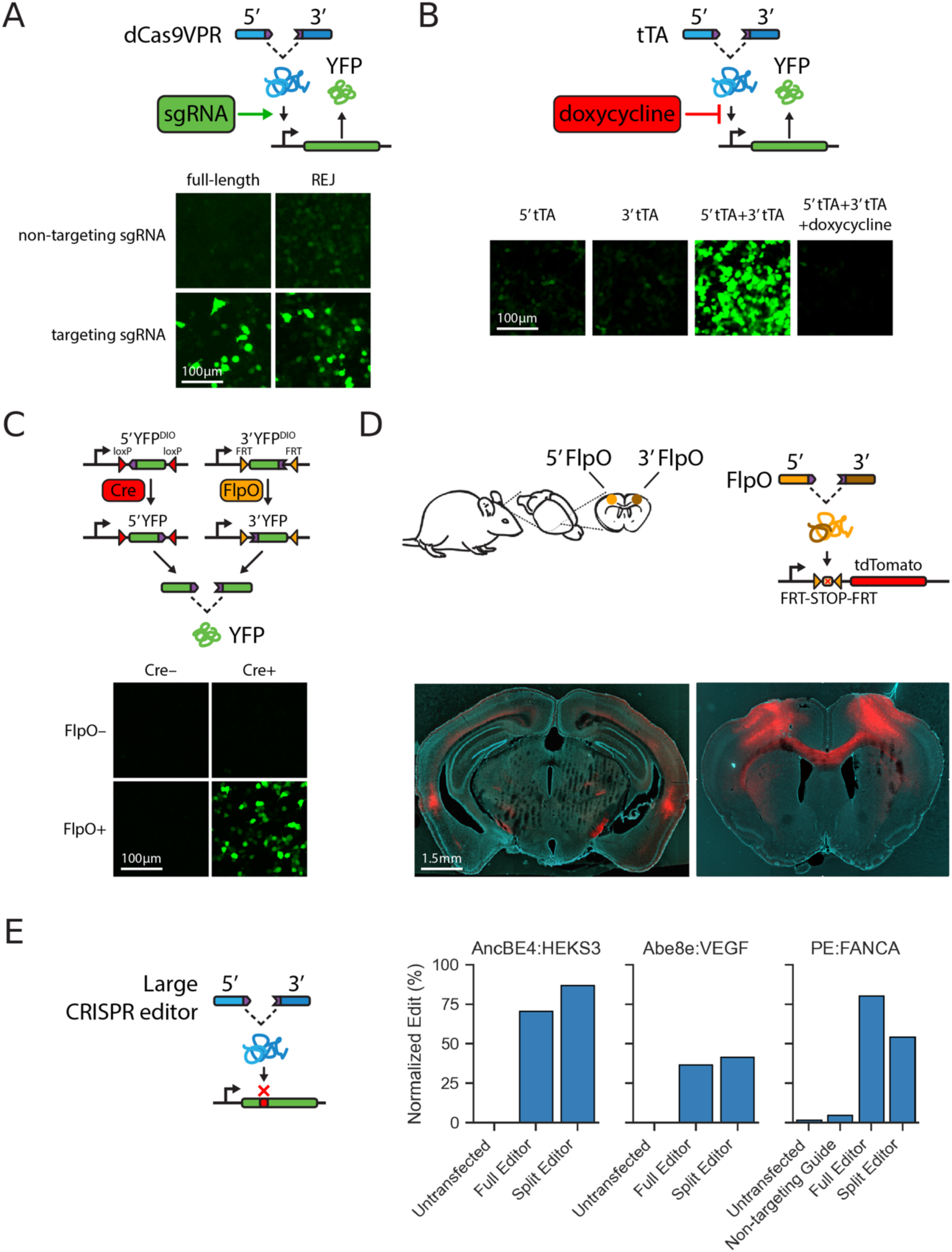
REJ enables the construction of synthetic gene circuit systems. **A)** REJ reconstitution of a split dCas9-VPR activator. HEK293T cells were transfected with a control construct encoding full-length dCas9-VPR or a split dCas9-VPR with REJ motifs. The addition of sgRNA allowed dCas9-VPR to activate a YFP reporter. **B)** Split tetracycline-controlled transcriptional activation (tTA) of YFP (Tet-OFF). When both 5’ and 3’ REJ sequences were delivered, the reconstituted tTA activated YFP expression and was repressible through the addition of doxycycline. **C)** REJ reconstitution of a split YFP using dual recombinase mediated intersectional genetics. HEK293T cells transfected with the Cre-dependent 5’ REJ RNA, Flp-dependent 3’ REJ RNA, a Cre expressing plasmid, and a FlpO expressing plasmid, resulted in successful reconstitution of YFP. **D)** REJ-enabled intersectional genetics with split FlpO encoded by dual AAVs. Following unilateral injection of each virus into a hemicortex only commissural neurons coexpress both the 5’FlpO and 3’FlpO REJ-RNAs. This reconstitutes FlpO activity which is detected by activation of a tdTomato reporter only in commissural neurons and labeling of their cell bodies and axons. **E)** Transfection-normalized editing efficiencies for full-length and split CRISPR editors across AncBE4:HEKS3, Abe8e:VEGF, and PE:FANCA show strong editing with split REJ designs (AncBE4:HEKS3, 87.1%; Abe8e:VEGF, 41.6%; PE:FANCA, 54.3%), demonstrating broad compatibility of REJ across distinct CRISPR effector platforms.

Many genetic studies have performed intersectional genetics using the co-expression of Cre and Flp to target specific cell types ^41^. We next tested whether the expression of a split-YFP reporter could be controlled by combinations of Cre and Flp (Figure 6C, Figure S3). We found that REJ-mediated expression of YFP could be exquisitely controlled by these recombinases, opening the possibility for creating multi-layer genetic circuits where the intersection of four separate components can be used to regulate transgene expression. This was further explored *in vivo* using a split-FlpO transgene to label callosal neurons within the cortex. Separate segments of REJ-5’ and REJ-3’ FlpO were injected into either the right or left cortex of a mouse strain carrying a FRT-tdTomato reporter (Figure 6D). This injection strategy co-infected only commissural neurons with the split-FlpO vectors, leading to selective activation of the tdTomato reporter in callosal projection neurons. Finally, we established that protein enzymatic activity could be efficiently reconstituted with REJ mediated trans-splicing by testing three different base editors: AncBE4, Abe8e, and Prime editor (PE). In each case the REJ-split editor performed with comparable efficiency to the full-length editor control (Figure 6E).

## Discussion

This report describes the optimization of short synthetic nucleotide modules that mediate RNA trans-splicing. This platform system, termed RNA end joining (REJ) to avoid confusion with non-RNA based trans-splicing ^42–45^, is functional across cell types and genes. The RNA reaction mediated by REJ engages the intrinsic cellular cis-splicing machinery to generate full length mRNAs that lack residual amino acids (scars) or mutations at the junction. We developed a set of intersectional genetic tools for biological studies and created an algorithm with user-weighted parameters for designing new gene expression systems with REJ. Importantly the system does not require extensive screening to achieve high expression levels of functional protein.

Furthermore, the REJ system is flexible and can be combined with different promoters, vectors and capsid types - thereby expanding its versatility. Below we describe the safety and efficiency parameters of REJ, and outline applications of the system for gene complementation, intersectional genetics, artificial genetic circuits and expression of large proteins.

### REJ Safety

There are several features of REJ that enhance safety. First, REJ-mediated RNA end-joining requires no foreign proteins that could trigger antigenic responses. Second, the system was designed to minimize translation of unspliced RNAs, with the goal of limiting the expression of protein fragments that may have a dominant negative activity. For example, gene therapy treatments that produce fragments of protein such as those based on inteins may necessitate higher dosing if dominant negative fragments attenuate the function of the therapeutic protein^46^. Third, the junctions between RNAs with complementary REJ modules were precise and the system is not prone to introducing frameshifts such as observed with ribozyme-based RNA joining ^36^. And fourth, the REJ system was optimized for efficiency by selecting efficient RDDs with particular stem-loop secondary structures that function well across a range of transcripts.

High efficiency is beneficial for lowering dosing levels that can trigger immune responses ^47^ and helps with viral manufacturing costs. We found dual vectors yield ∼40%-60% of the protein levels produced by single vector controls across a range of different transcripts. In general, the highest efficiency for expression of any particular gene with REJ was achieved when equimolar levels of each RNA were co-expressed. In samples where both RNA and protein levels were monitored, we found that REJ-spliced transcripts were translated with higher efficiency than control transcripts lacking introns. Possibly REJ-mediated trans-splicing, like normal intronic cis-splicing, enhances the transport, stability, and/or translation of mRNAs ^29^.

To examine whether REJ transcripts are inappropriately trans-spliced to endogenous cellular RNAs, we infected mouse muscle with dual REJ-AAV vectors and performed RNA sequencing. Endogenous trans-splicing was not detected *in vivo*, whereas in extreme conditions in HEK 293T cells and with deep sequencing, off-target interactions were only observed with the 3’ REJ-RNA at very low levels that may be at a frequency of less than 1 event per cell. Taken together, these findings suggest that under typical conditions, REJ off-target splicing to cellular RNAs is unlikely to be deleterious.

An additional safety concern with the REJ system is that it is mediated by RNA::RNA interactions that could trigger intrinsic cellular dsRNA sensors for antiviral responses^38,48^. The RNA::RNA pairs formed by the RDDs of REJ function in the nucleus to promote splicing, which is predicted to reduce the level of dsRNA in the cytoplasm where the sensors are located^49^.

Consistent with this, we did not observe overt changes in cell growth, survival or protein expression associated with REJ-mediated gene expression *in vitro* or *in vivo*. Likewise, transcriptomic analysis from cells expressing REJ constructs did not detect dsRNA antiviral responses. Possibly the length or concentration of cytoplasmic REJ-dsRNA is below the threshold for detectors such as RIG-I and MDA5^50–54^.

### REJ and Other Dual Vector Systems

AAV has a small 25nm particle size, which facilitates the broad spread of this vector within tissues. However, this size limits the cargo capacity of single strand AAV to ∼4.5kb, and ∼2.5kb for faster-replicating self-complementary AAVs. Many genes exceed these cargo limits, particularly when combined with tissue specific promoters. Because multiple AAVs can readily coinfect cells, several dual vector approaches have emerged to address these cargo constraints^36,37,44,55^. Dual vector systems are based on splitting genes among multiple viruses, which coinfect cells and recombine the genes at the level of DNA^32^, RNA (Figure 7A)^18^, or protein^43,44,56^. We found that the reconstitution of split-YFP was 15-fold greater with REJ compared to DNA recombination.

**Figure 7:**
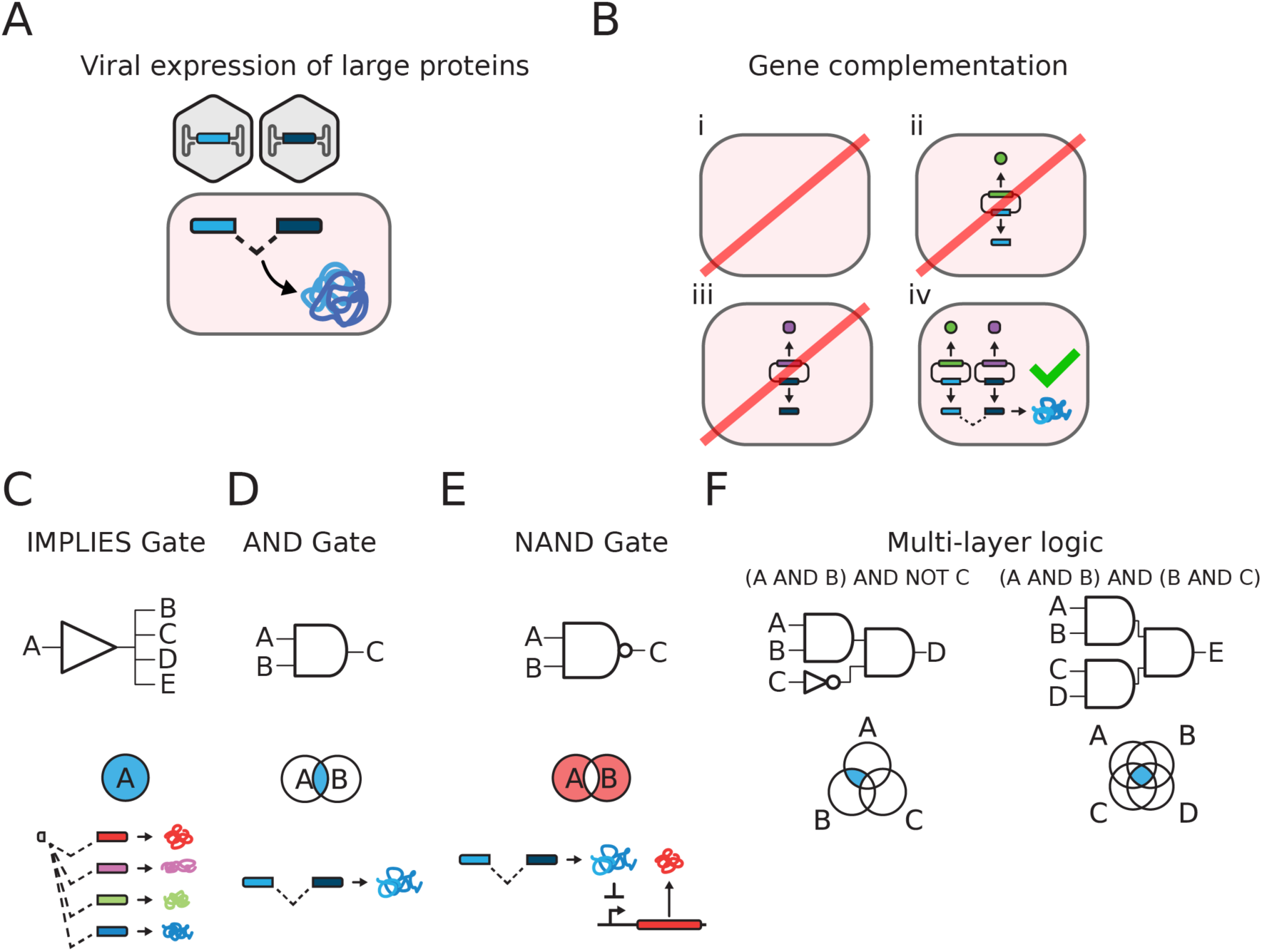
REJ enables broad biological applications. **A)** REJ mediated trans-splicing can facilitate the viral expression of large proteins, expanding payload capacity beyond single-AAV packaging limits. **B)** REJ mediated trans-splicing can enable intersectional selection with a single split marker gene (blue). Only cells receiving both constructs survive, with each construct able to carry additional cargo (green, purple). **C-F)** REJ-mediated complementation for creating advanced logic circuits for targeted gene expression. **C)** Schematic of a REJ mediated IMPLIES circuit with a constant 5’ leader REJ RNA used to join with multiple 3’ REJ RNAs to co-express multiple protein isoforms (e.g. see Figure S4). In principle the ratio of 3’ RNAs could be manipulated to control the relative level of each isoform. **D)** Schematic of a REJ-mediated AND circuit where both REJ RNAs are conditionally expressed in the same cell thereby allowing intersectional protein expression (e.g. see Figure 6D). **E)** Schematic of a REJ mediated NAND circuit in which coexpression of both REJ RNAs leads expression of a transcriptional repressor or cell death toxin (e.g. TetOFF, see Table 1). **F)** Schematic of two multi-layer logic systems with REJ, allowing for complex intersectional conditional gene expression. An (A AND B) AND NOT C circuit can be implemented with a split Tet-OFF system where A, B, and C are 5’ REJ RNA, 3’ REJ RNA, and doxycycline respectively (e.g. see Figure 6B). An AND(A, B, C, D) circuit is demonstrated as a dual recombinase intersectional genetic system (e.g. see Figure 6C).

Inteins that mediate “protein splicing” have proven to be an effective means for expressing efficacious levels of protein from dual AAVs ^43,44,56^. Inteins range in size from ∼100-800aa and can be appended to transgenic proteins to catalyze the joining of separate fragments. There are, however, important considerations when using inteins. Following intein catalysis a ∼3-6aa residual scar remains at the junction site. Therefore, it is necessary to screen whether the insertion compromises transgenic protein function. In addition, inteins are derived from bacteria, yeast and archaea and may therefore be detected as foreign epitopes that trigger immune responses in mammals. Finally, the protein fragments that are the substrate for intein-joining may have dominant negative activity since they are truncations of the full length protein.

The use of a self-inactivating Cre system termed AAVLINK for joining dual-AAV vectors at the level of DNA is a promising strategy for expressing large proteins ^37^, however it relies upon a potentially-antigenic recombinase from bacteriophage that may have antigenic and cytotoxic activity ^57^. Thus, the expression of large proteins using RNA-based solutions could provide an added level of safety. In this regard, dual vector systems based on RNA trans-splicing have been previously described ^18,19,31,58^. The REJ system developed here is an extension of these RNA trans-splicing systems. We optimized REJ modules so that RNA trans-splicing can be utilized across a wide range of transcripts without the need for large-scale screening. The REJ modules mediate precise and efficient RNA trans-splicing, and include design features that limit expression of protein fragments encoded by unspliced mRNAs. We generated a battery of split reporter genes, developed tools for joining ≥2 RNAs, and provide a computational algorithm and web tool that facilitates the design of efficient trans-splicing systems.

An alternative to coopting cellular splicing machinery for RNA trans-splicing is the development of ribozyme-based approaches such as STITCHR ^36^. While this is a very promising technology, the mechanistic features that control the reactions between exposed hydroxyl and cyclic-phosphate groups from ribozyme cleavage are not yet well understood and therefore the variables that impact safety, off-target reactions, and efficiency in the context of different cells and transcripts are uncertain. In many ways ribozyme and trans-splicing systems have similar advantages, however the REJ system described here can be engineered to join ≥2 RNA fragments for more complex applications. Multiple dual vector systems may prove to have clinical value for treating genetic diseases caused by loss-of-function mutations in large genes, however this general approach does add complexity for manufacturing and safety testing that may one day be superseded by nanoparticle delivery systems ^59^.

### Creating Synthetic Gene Regulation Circuits with REJ

Genetic complementation has been extensively used to study protein function and cell selection (Figure 7B)^60^. For example, AND gate genetic systems are convenient for identifying cells that have been cotransfected with multiple constructs. The REJ system can be used to select cotransfected cells with a single marker by including a split fluorescent reporter or antibiotic resistance gene on each plasmid (Figure 7B, E; Table 1). Likewise, intersectional genetics is often used to target discrete cell types *in vivo*. The REJ system can be used to simplify complex combinatorial labeling using the expression of a single reporter (Figure 6D, 7E).

Building on the use of combinatorial genetics to create gene circuits, we found that IMPLIES gates could be created by combining a 5’ REJ-RNA with a mixture of different 3’ REJ-RNAs, leading to the co-expression of multiple protein isoforms within cells (Figure S4, Figure 7C). Likewise, NAND and Multi-layer logic gates can be created using intersectional complementation strategies with regulatory proteins such as TetOFF and dCas9-VPR systems (Figure 6A-C, 7D-G; Table 1). Taken together, the REJ system can be used as a versatile gene expression platform for creating novel gene circuits that facilitate the study of specific cell types as well as open additional possibilities for gene therapy.

## Methods

### RESOURCE AVAILABILITY

#### Lead contact

Further information and requests for resources and reagents should be directed to and will be fulfilled by the lead contact, Samuel L. Pfaff.

#### Materials availability

Plasmids generated for this study, including REJ toolkit constructs, are deposited to Addgene. Plasmids and other unique reagents that are not available through Addgene are available from the lead contact upon reasonable request.

#### Data and code availability

High-throughput sequencing data generated in this study have been deposited at the NCBI Sequence Read Archive (SRA) and are publicly available as of the date of publication. Targeted and deep-sequencing data from HEK293T cell experiments (REJ splicing-fidelity and endogenous off-target analyses) are available under NCBI BioProject accession PRJNA1480053. RNA-sequencing data from AAV-treated mouse muscle (whole-transcriptome safety profiling and *in vivo* off-target splicing analyses) are available under NCBI BioProject accession PRJNA1480054. Original code for the REJenerator design algorithm and the REJ Studio web application has been deposited on GitHub and is publicly available (https://github.com/rlfarman/rej-studio). The REJenerator algorithm and the transcriptome-scale resource of REJ-ready coding sequences are accessible through the REJ Studio web application at www.rejstudio.com. During peer review the site is access-restricted; reviewers may log in using a password. The site and the REJ-ready sequence resource will be made publicly available without restriction upon publication. Any additional information required to reanalyze the data reported in this paper is available from the lead contact upon request.

#### Cell culture models

HEK293T cells were maintained in 1X Dulbecco’s Modified Eagle Medium (DMEM; Gibco) supplemented with 10% fetal bovine serum at 37 degrees C and 5% CO2. HEK293T cells were used for all in vitro plasmid transfection experiments in this study.

#### Mouse models and husbandry

All animal procedures were performed in accordance with the NIH Guide for the Care and Use of Laboratory Animals and approved by the Salk Institute for Biological Studies Institutional Animal Care and Use Committee (IACUC protocol 23-00020). C57BL/6J mice of both sexes were used for in vivo experiments. Mice were housed in a specific pathogen-free facility under a 12-hour light/dark cycle with ad libitum access to standard chow and water.

### METHOD DETAILS

#### REJ construct design and nomenclature

REJ constructs were designed so that separate transcriptional units encode portions of a target coding sequence that are reconstituted by RNA trans-splicing. For dual-vector designs, the upstream fragment was designated the 5’ REJ RNA and contained the 5’ portion of the coding sequence followed by a synthetic REJ module, whereas the downstream fragment was designated the 3’ REJ RNA and contained a corresponding REJ module followed by the 3’ portion of the coding sequence. For triple-vector designs, coding sequences were divided into 5’, middle (M), and 3’ REJ RNAs. REJ modules contained engineered RNA dimerization domains and splice cis-elements positioned to promote spliceosome-mediated RNA trans-splicing and removal of the REJ module sequences from the mature, scar-free reconstituted mRNA.

#### RNA dimerization domain design

RNA dimerization domains (RDDs) were designed to promote sequence-specific RNA:RNA interactions between REJ RNAs. Three general RDD architectures were evaluated: linear (Lin), hypodiverse loop (HDL), and kissing stem loop (KSL) designs. Lin RDDs consisted of 50-mer random nucleotide sequences encoded in reverse-complement orientation between the 5’ and 3’ REJ RNAs. KSL RDDs contained three tandem hairpin loops separated by 4-uracil spacers; each hairpin contained a 10-bp stem and a 10-nt loop. Matched RDD pairs were tested for their ability to promote REJ-mediated split-YFP reconstitution, whereas mismatched or non-pairing RDD combinations were used to assess pairing specificity. Orthogonal RDD pairs were also evaluated to support multipart REJ designs in which more than two RNA segments must be joined in a defined order.

#### REJ module engineering and cis-element mutagenesis

REJ module variants were generated to test the contribution of splice cis-elements and expression-enhancing features to RNA trans-splicing efficiency. Splice donor and splice acceptor motifs were disrupted by scrambling each motif while maintaining its original nucleotide composition, and these variants were used to test whether split-YFP reconstitution required canonical splice sites. Intronic splice enhancer elements, including the 5’ DISE, 5’ ISE, and 3’ ISE, were deleted individually or in combination to assess their contribution to REJ activity. Additional construct variants were generated to evaluate the effects of WPRE inclusion in the 3’ REJ construct and stimulatory cis-introns on reporter expression.

#### Split-cargo and split-site selection

Genetic cargos used in this study were selected to evaluate REJ across the applications described in the Results, including reporter, therapeutic, genome-editing, transcriptional-regulatory, recombinase, and selection-marker contexts. For each cargo, the coding sequence was manually divided into two segments, with candidate split sites chosen approximately near the midpoint of the coding sequence. Split sites were selected at, or synonymously recoded to create, a WG|GW motif while preserving the encoded protein sequence. Because REJ reconstitutes the mature coding sequence through scar-free RNA trans-splicing, split-site selection did not require the unspliced RNA fragments to encode productive in-frame protein segments. Coding sequences were codon optimized for human expression during construct design.

#### Plasmid assembly, bacterial propagation, and sequence validation

Plasmids were designed using SnapGene software and assembled using In-Fusion HD (Takara) or NEBuilder HiFi (NEB) kits. REJ constructs were generated from designed coding sequences, REJ modules, promoters, fluorescent transfection markers, and other regulatory elements as required for each experiment. Assembled plasmids were transformed into Stellar Competent Cells (Takara) and propagated under antibiotic selection. Final plasmids were purified for transfection or viral packaging using QIAprep Mini (Qiagen), ZymoPURE Midi (Zymo), or PureLink Maxi (Invitrogen) kits. Plasmid identities were verified by control restriction digests using NEB restriction enzymes and by Sanger sequencing across assembled junctions and engineered regions.

#### Cell culture and plasmid transfection

HEK293T cells were maintained in DMEM (Gibco) supplemented with 10% FBS at 37 degrees C and 5% CO2. For plasmid transfection experiments, cells were seeded in 48-well plates and transfected at approximately 80% confluency with 600 ng of each indicated plasmid using Lipofectamine 3000 according to the manufacturer’s protocol. Unless otherwise indicated, cells received the plasmids specified for each experiment, including 5’ REJ RNA, 3’ REJ RNA, full-length control, reporter, guide RNA, recombinase, or transfection-marker plasmids as appropriate. For experiments comparing 5’ and 3’ REJ RNA combinations, plasmids were co-transfected at the indicated stoichiometric ratios. Cells were analyzed or harvested 48 hours after transfection for fluorescence imaging, reporter quantification, RNA extraction, or protein extraction.

#### Split-YFP reporter imaging and quantification

Split-YFP reporter assays were performed in HEK293T cells transfected with plasmids encoding the indicated 5’ REJ RNA and 3’ REJ RNA constructs. Where indicated, the 5’ and 3’ REJ plasmids carried BFP and RFP transfection markers, respectively, and full-length YFP controls were transfected with a single fluorescent transfection-control marker. For Figure 1C, cells were imaged 48 hours after transfection using an Olympus FV3000-RS microscope, segmented using the Cellpose 3 cyto model, and YFP fluorescence was quantified from segmented cells. For all other split-YFP reporter quantification experiments, cells were analyzed by flow cytometry. Dual-transfected cells were identified by gating on BFP and RFP marker expression as appropriate, and YFP levels were quantified within the dual-positive population. Full-length YFP control samples were analyzed using the corresponding single transfection-control marker. YFP signal was compared with the relevant full-length YFP or matched control condition as indicated in the figure legends.

#### Fluorescence reporter assays for REJ architecture screens

REJ architecture variants in Figures 1 and 2 were evaluated using split-YFP fluorescence reporter assays in HEK293T cells. Matched, mismatched, or non-pairing RDD combinations were tested by co-transfecting the corresponding 5’ REJ RNA and 3’ REJ RNA plasmids and measuring YFP reconstitution by flow cytometry in marker-positive cells. Orthogonal RDD pairs were evaluated by comparing matched and mismatched pairings, and triple-vector YFP designs were tested by co-transfecting 5’, middle (M), and 3’ REJ RNA plasmids. REJ cis-element variants were tested using the same split-YFP reporter framework, including splice donor and splice acceptor scrambling, deletion of intronic splice enhancer elements, WPRE inclusion, and addition of stimulatory cis-introns. Reporter output was normalized to the indicated matched control or full-length YFP condition for each experiment.

#### Protein extraction and western blot analysis

For western blot experiments, HEK293T cells were harvested 48 hours after transfection, washed with PBS, and lysed in RIPA lysis buffer (Cell Signaling Technology, Cat#9806) supplemented with EDTA-free protease inhibitor (Sigma, Cat#11873580001) and Benzonase. Lysates were passed through an insulin syringe three times and centrifuged for 10 minutes at 21,300 rcf. Supernatants were collected and stored at −20 degrees C until analysis. Protein concentration was measured using the Pierce BCA Protein Assay Kit (Thermo Scientific, Cat#23227). Samples were separated on NuPAGE 4-12% Bis-Tris gels (Thermo Fisher, Cat#NP0321BOX or Cat#NP0323BOX) or NuPAGE 7% Tris-Acetate gels (Thermo Fisher, Cat#EA03585BOX) using NuPAGE MOPS running buffer (Thermo Fisher, Cat#NP0001) or Tris-Acetate running buffer (Thermo Fisher, Cat#LA0041), as appropriate for the target protein size. Proteins were transferred using NuPAGE transfer buffer (Thermo Fisher, Cat#NP00061) onto Immobilon-FL transfer membranes (Millipore Sigma, Cat#IPFL00010). Membranes were blocked in PBS/casein/Tween buffer, washed in PBS with 0.1% Tween20 (Sigma-Aldrich, Cat#P7949), and probed with anti-HA or anti-FLAG primary antibodies. Fluorescent secondary antibodies were used for detection, and blots were imaged and quantified using ImageStudio software.

#### ABCA4 REJ and intein comparison

ABCA4 expression from dual-vector REJ and dual-vector intein designs was compared in HEK293T cells by plasmid transfection followed by western blot analysis. Cells were transfected with plasmids encoding full-length ABCA4, the paired ABCA4 REJ constructs, or the paired ABCA4 intein constructs. Protein lysates were collected 48 hours after transfection and analyzed as described above. Full-length ABCA4 signal and C-terminal fragment band signal were quantified from western blot images using ImageStudio. Product purity was assessed by comparing the abundance of full-length ABCA4 with detectable C-terminal fragment bands, and REJ and intein conditions were normalized to the full-length ABCA4 control as indicated in the figure legends.

#### CRISPR editor and transcriptional activator activity assays

REJ compatibility with CRISPR-based effectors was tested in HEK293T cells by comparing full-length effector plasmids with paired 5’ REJ RNA and 3’ REJ RNA split-effector plasmids. For dCas9-VPR transcriptional activation assays, cells were co-transfected with full-length or split dCas9-VPR constructs, the indicated sgRNA, and a YFP reporter plasmid. YFP activation was assessed 48 hours after transfection by fluorescence microscopy using the Olympus FV3000-RS microscope, with non-targeting sgRNA and reporter-only conditions used as negative controls.

For genome-editing assays, full-length or split REJ editor constructs were co-transfected with the corresponding guide RNA and target components for each editor system. Editing activity at the indicated target sites for AncBE4:HEKS3, Abe8e:VEGF, and prime editor:FANCA was measured by targeted amplicon next-generation sequencing. Editing percentages were quantified using CRISPResso2, and split-editor activity was compared with the corresponding full-length editor control.

#### Tet and recombinase-based intersectional reporter assays

REJ-based intersectional reporter assays were performed in HEK293T cells using plasmid combinations specified for each circuit. For tetracycline-regulated reporter assays, cells were transfected with split REJ constructs encoding Tet-OFF tTA together with the corresponding Tet-responsive YFP reporter plasmid. Component omission controls were included to test whether reporter activation depended on the expected combination of inputs. YFP expression was assessed 48 hours after transfection by fluorescence microscopy using the Olympus FV3000-RS microscope.

For recombinase-based intersectional assays, cells were transfected with lox- and FRT-inverted 5’ REJ RNA and 3’ REJ RNA coding sequences together with Cre and FlpO expression plasmids as indicated. Reconstitution of YFP was assessed by fluorescence microscopy using the Olympus FV3000-RS microscope, and component omission controls were used to verify intersectional dependence.

#### Recombinant AAV production, purification, and titration

Recombinant AAV vectors were produced in HEK293T cells by triple transfection with the relevant Rep/Cap plasmid, Ad5 helper plasmid, and transfer plasmid by the Salk Viral Vector Core. AAV8 or AAV9 capsids were used as indicated for each experiment. Viral particles were harvested 72 hours after transfection by three freeze-thaw cycles and Benzonase treatment, followed by PEG 8000 precipitation. Vectors were purified using iodixanol gradients or sequential CsCl density gradients, buffer-exchanged into PBS with 5% sorbitol, sterile-filtered, and titered by qPCR in triplicate.

#### In vivo AAV delivery

For in vivo REJ-YFP experiments, AAV vectors encoding the indicated 5’ REJ RNA and 3’ REJ RNA constructs were mixed before delivery. Dual-vector REJ conditions received matched 5’ and 3’ REJ AAVs, and single-vector YFP controls received an AAV encoding full-length YFP. Mice were anesthetized using a blend of fentanyl (0.05 mg/kg), midazolam (5 mg/kg), and medetomidine/Dormitor (0.5 mg/kg). For intramuscular delivery, vectors were injected into the tibialis anterior muscle. For systemic delivery, AAV vectors were administered by retro-orbital intravenous injection. For cortical delivery, AAV vectors were injected stereotaxically into the brain at coordinates defined relative to bregma; the anterior-posterior coordinate was −2.54 mm from bregma. For retinal delivery, AAV vectors were delivered by subretinal injection. Viral dose, capsid, injection route, stereotaxic coordinates, and collection time point are reported for each experiment where available.

#### Tissue collection, fixation, sectioning, and native fluorescence preparation

Tissues were collected at the indicated experimental endpoints after AAV delivery. Dissected tissues were fixed by immersion in 4% paraformaldehyde (Electron Microscopy Sciences, Cat#15713), washed in PBS, cryoprotected in PBS containing 30% sucrose at 4 degrees C, embedded in OCT (Tissue-Tek, Cat#4583), frozen, and cryosectioned onto glass slides (Fisherbrand, Cat#12-550-15). Tissue sections were prepared for YFP fluorescence imaging without antibody staining. Hoechst nuclear stain (Invitrogen, Cat#H3570) was used for nuclear visualization where shown. Sections were mounted in Mowiol and imaged on an Olympus FV3000-RS confocal microscope. Macroscopic YFP fluorescence from dissected tissues was imaged using a the Zeiss Lumar.V12 fluorescence stereomicroscope. Imaging settings were kept consistent within each experiment when comparing control and REJ-treated conditions, and YFP fluorescence was quantified from images where indicated.

#### RNA extraction, RT-PCR/qPCR, and junction-spanning assays

Total RNA was isolated from cells or tissues using the RNeasy Plus Universal Mini Kit (Qiagen, Cat#1062832) and treated with RNase-Free DNase Set (Qiagen). RNA isolates were reverse transcribed to cDNA using AccuPower RT PreMix (Bioneer). Junction-spanning qPCR was performed in technical triplicate using primers positioned on opposite sides of the expected REJ splice junction. For YFP reconstitution assays, junction-spanning qPCR was used to compare split-REJ YFP transcript abundance with full-length YFP transcript abundance. REJ junction amplicons generated for sequence validation were analyzed by Sanger sequencing as described below.

#### Sanger sequencing of REJ splice junctions

To validate REJ splice-junction fidelity, cDNA generated from cells expressing paired REJ constructs was amplified using primers flanking the expected mature splice junction. PCR products were purified using the DNA Clean & Concentrator-5 kit (Zymo Research, Cat#D4003) and submitted to Eton Bioscience for Sanger sequencing. Sequencing chromatograms were inspected across the REJ junction to confirm precise, scar-free joining at the expected splice boundary.

#### RNA-seq library preparation and sequencing

Total RNA was isolated from AAV-treated muscle tissue using the RNeasy Plus Universal Mini Kit (Qiagen, Cat#1062832). RNA integrity was assessed using an Agilent Bioanalyzer 4200. Library preparation and sequencing were performed by the Salk Next-Generation Sequencing Core. Libraries were prepared using the Illumina TruSeq Stranded mRNA kit with polyA enrichment and sequenced on an Illumina NovaSeq 6000 SP with 150 bp paired-end reads at a target depth of 25 million reads per sample. Raw reads were quality controlled and trimmed using USEARCH12. Transcript abundance was quantified by mapping reads to the GRCm39/mm39 mouse reference transcriptome using Salmon.

#### REJ splicing-fidelity and endogenous off-target analysis

REJ splicing fidelity and endogenous off-target splicing were quantified from RNA-seq reads using custom Python scripts. Reads were filtered for 25-bp anchor sequences immediately upstream or downstream of the expected REJ splice junction in the 5’ REJ RNA or 3’ REJ RNA. The adjacent 15-bp sequence next to each anchor was extracted and classified as on-target spliced, unspliced, or candidate endogenously spliced based on whether it matched the expected partner REJ segment, retained the unspliced REJ module sequence, or matched transcriptome sequence. For each REJ RNA, classified reads were counted and expressed as the proportion of total classified reads. Endogenous off-target rates and upper confidence bounds were calculated from the number of observed endogenous off-target splice products.

#### Whole-transcriptome safety and innate immune-response analysis

Whole-transcriptome RNA-seq analysis was used to compare muscle samples from saline-treated, single-AAV YFP-treated, and dual-AAV REJ-YFP-treated mice. Transcript abundance quantification generated with Salmon was analyzed for sample-level structure by principal component analysis using scikit-learn. Differential gene expression analysis was performed using PyDESeq2 for pairwise comparisons among treatment groups. Genes were considered differentially expressed using the significance and fold-change thresholds indicated in the figure legends. Innate immune-response and double-stranded RNA sensor pathway activation were assessed from the muscle RNA-seq data by examining expression of relevant antiviral and pattern-recognition-response genes.

#### REJenerator design algorithm

REJenerator was implemented in Python using DNA Chisel to automate design of REJ-compatible coding sequences. Starting from an input coding sequence, the algorithm performs synonymous recoding while preserving the encoded amino acid sequence. Sequence optimization simultaneously reduces cryptic splice donor and splice acceptor motifs, reduces CpG dinucleotide content, controls k-mer complexity, maintains GC content between 35% and 60%, and improves codon adaptation index (CAI) for the selected species. After coding-sequence optimization, REJenerator scans the user-specified split region to identify or introduce a suitable WG|GW junction motif near the target split coordinate and outputs the corresponding 5’ REJ RNA and 3’ REJ RNA coding sequences for construct generation. REJenerator is accessible through the REJ Studio webtool at www.rejstudio.com (during peer review, a password is required to log in), and the source code for the REJenerator algorithm and REJ Studio web application is available at https://github.com/rlfarman/rej-studio.

#### Transcriptome-scale generation of REJ-ready coding sequences

An empirical WG|GW motif analysis was performed to determine how readily REJ-compatible splice junction motifs could be identified or introduced by synonymous recoding. A total of 10,000 coding sequences were randomly sampled from the human transcriptome, and each sequence was analyzed to determine the distance from candidate split coordinates to the nearest WG|GW motif that could be induced while preserving the encoded protein sequence.

REJenerator was then applied transcriptome-wide to all human and mouse protein isoforms longer than 65 amino acids to generate REJ-ready coding sequences as a public resource. For each coding sequence, REJenerator optimized synonymous sequence features as described above and recorded splice donor counts, splice acceptor counts, CpG dinucleotide counts, k-mer complexity, GC content, and CAI before and after optimization. Coding sequences were stratified by length to compare genes that do not explicitly require multi-AAV delivery (<4,000 bp), genes requiring a two-part multi-AAV design (>=4,000 and <8,000 bp), and genes requiring a three-part multi-AAV design (>=8,000 bp). The resulting REJ-ready sequence resource will be made publicly available.

#### Data visualization and figure generation

Quantitative data were analyzed and plotted in Python using seaborn and matplotlib. Flow cytometry data were analyzed using FlowJo software. Fluorescence microscopy images were processed using ImageJ/Fiji, Cellpose 3, or Python-based analysis scripts as indicated. Western blot images were quantified using ImageStudio. Final figure panels were assembled in Adobe Illustrator.

## QUANTIFICATION AND STATISTICAL ANALYSIS

General statistical analysis

Statistical analyses were performed in Python using scipy. Data are presented as mean +/-SEM unless otherwise specified. Pairwise comparisons were performed using Welch’s t test. Sample sizes, replicate definitions, and exact p-values are reported in the figure legends.

## Acknowledgements

S.L.P. is the Benjamin H. Lewis Chair in Neuroscience. This research was supported by the Howard Hughes Medical Institute, DoD MD190071, National Institute of Neurological Disorders and Stroke (NINDS) 1 RO1 NS123160-01, the Usher 1F Foundation, Salk Institute Gene Therapy Fund, Keck Foundation, Mathers Foundation, and the Sol Goldman Charitable Trust. R.H.H. was supported by F30 HD106732. The Salk NGS Core is supported with funding from NIH-NCI CCSG: P30 014195. We thank Kathryn Hilde, Kenneth Harrison and Joe Markson for valuable comments on the research.

## Author Contributions

Conceptualization, L.C.B., R.H.H., K.J.H., C.E.W., S.K., and S.L.P.; Methodology, L.C.B., R.H.H., K.J.H., C.E.W., S.K., and S.L.P.; Software, L.C.B., R.H.H., R.L.F., and S.K.; Validation, L.C.B., R.H.H., K.J.H., C.E.W., and S.L.P.; Formal Analysis, L.C.B., R.H.H., K.J.H., and S.K.; Investigation, L.C.B., R.H.H., K.J.H., C.E.W., N.C., C.C., S.K., C.T.P., S.R., and K.L.; Resources, L.C.B., K.J.H., C.E.W., R.L.F., S.K., K.L., and S.L.P.; Data Curation, L.C.B., K.J.H., and C.E.W.; Writing – Original Draft, L.C.B., R.H.H., C.E.W., S.K., and S.L.P.; Writing, Review & Editing, L.C.B., R.H.H., C.E.W., S.K., and S.L.P.; Visualization, L.C.B., R.H.H., and K.J.H.; Supervision, L.C.B. and S.L.P.; Project Administration, L.C.B. and S.L.P.; Funding Acquisition, L.C.B., S.K., and S.L.P.

## Declaration of Interests

The Salk Institute for Biological Studies holds two patents on the RNA end-joining (REJ) technology described in this work, both of which have been licensed to Insmed. S.L.P. and L.C.B. are inventors on a patent covering the core REJ technology, and R.H.H., C.E.W., K.J.H., L.C.B., and S.L.P. are inventors on a second patent covering additional REJ technology. L.C.B. is an employee of Insmed, and R.H.H. is a consultant for Insmed. The remaining authors declare no competing interests.

## Declaration of generative AI and AI-assisted technologies in the writing process

During the preparation of this work, the authors used Claude (Anthropic) and ChatGPT (OpenAI) in order to edit and condense text for clarity. After using these tools, the authors reviewed and edited the content as needed and take full responsibility for the content of the published article.

---

**Figure S1.**
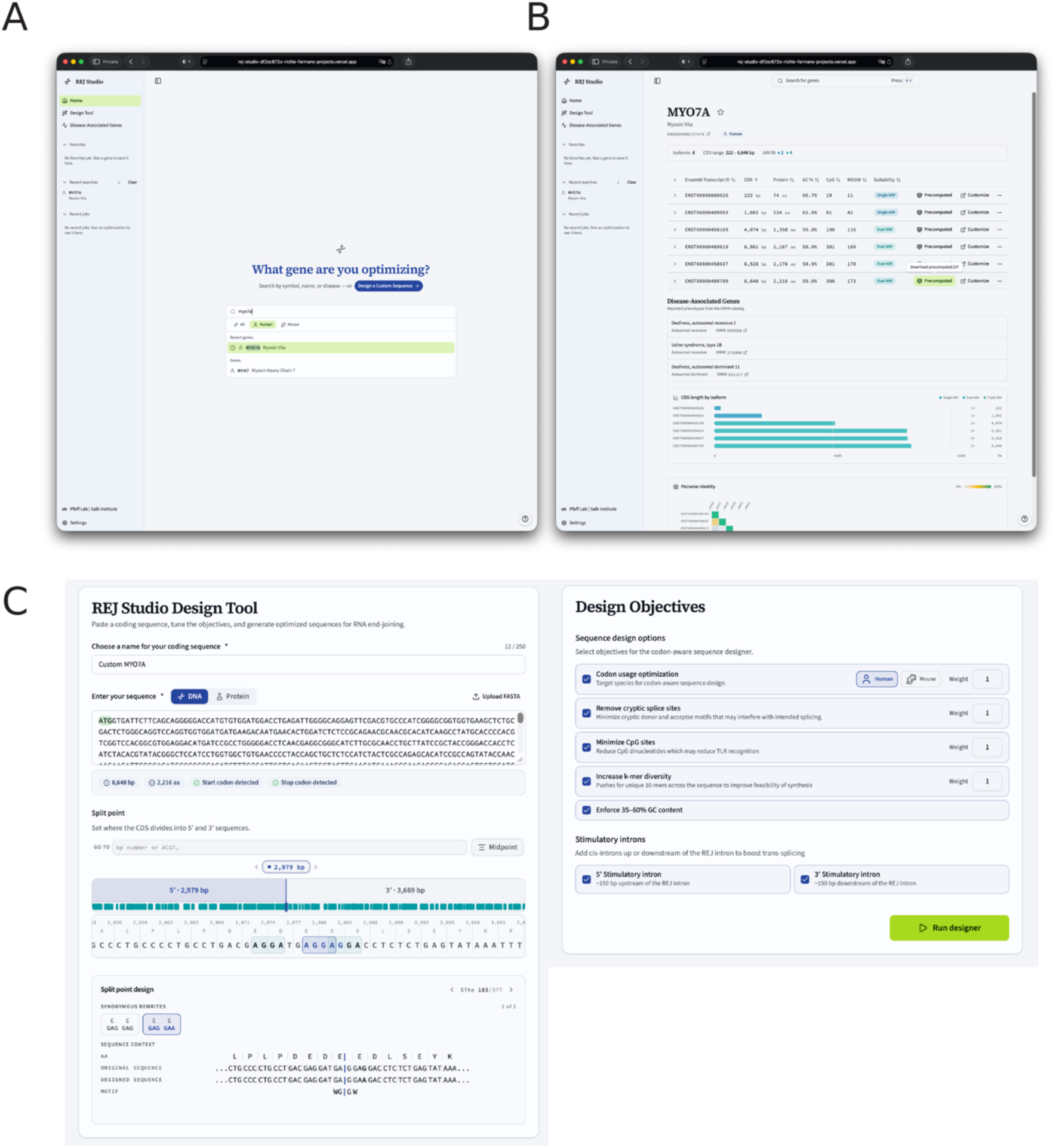
REJ Studio web interface for identifying and designing REJ-compatible split-gene constructs. (A) Landing page of REJ Studio. Users can search for any gene of interest by name to access precomputed REJ designs. The left sidebar provides navigation to launch the custom-sequence design tool or browse large disease-associated genes whose coding sequences exceed the packaging capacity of a single AAV vector. (B) Example search results page for MYO7A. The query returns annotated isoforms for the selected gene, illustrating how REJ Studio organizes transcript-specific design outputs. Precomputed REJ-ready sequences can be downloaded directly from the search results page, or imported into the design tool for further customization. (C) Custom REJ design interface. Users input a coding sequence and define the nucleotide region to inspect using a draggable sequence-position slider. Compatible split motifs within the selected region can be inspected, and synonymous codon substitutions can be selected to install a WG|GW splice-junction motif at the chosen position. A splice-junction preview displays the resulting sequence context. Additional design parameters allow users to enable or disable, and specify relative weights for, species-specific codon optimization, depletion of cryptic splice sites, CpG minimization, increased k-mer diversity, GC-content constraints, and induction of 5′ and 3′ stimulatory intron sites. Running the designer generates a downloadable zip file containing REJ-optimized sequences produced with the selected split-site and design parameters.

**Figure S2.**
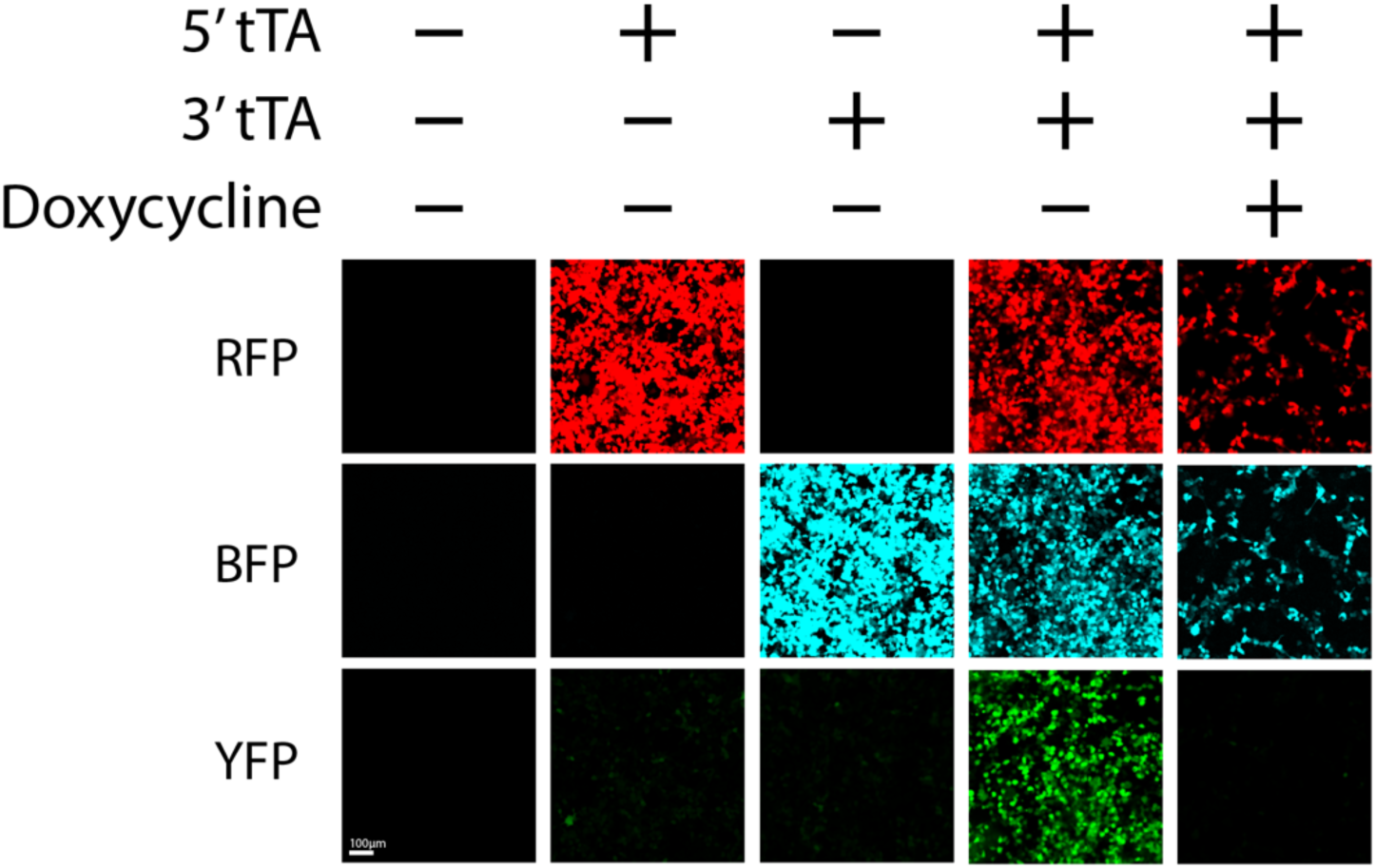
Combinatorial control of Tet-OFF reporter activation by REJ-mediated tTA reconstitution. Representative fluorescence images of HEK293T cells transfected with combinations of 5′ REJ-tTA and 3′ REJ-tTA constructs, with or without doxycycline, to test component-dependent activation of a Tet-responsive YFP reporter. All conditions included the Tet-responsive YFP reporter plasmid. Columns show cells receiving no REJ-tTA components, 5′ REJ-tTA alone, 3′ REJ-tTA alone, both 5′ REJ-tTA and 3′ REJ-tTA, or both REJ-tTA components plus doxycycline. Rows show fluorescence from the RFP marker co-expressed with the 5′ REJ-tTA construct, the BFP marker co-expressed with the 3′ REJ-tTA construct, and YFP reporter output. YFP expression was detected only when both 5′ REJ-tTA and 3′ REJ-tTA were present, consistent with REJ-mediated reconstitution of functional tTA. Addition of doxycycline suppressed YFP expression, confirming doxycycline-dependent inhibition of the Tet-OFF circuit.

**Figure S3.**
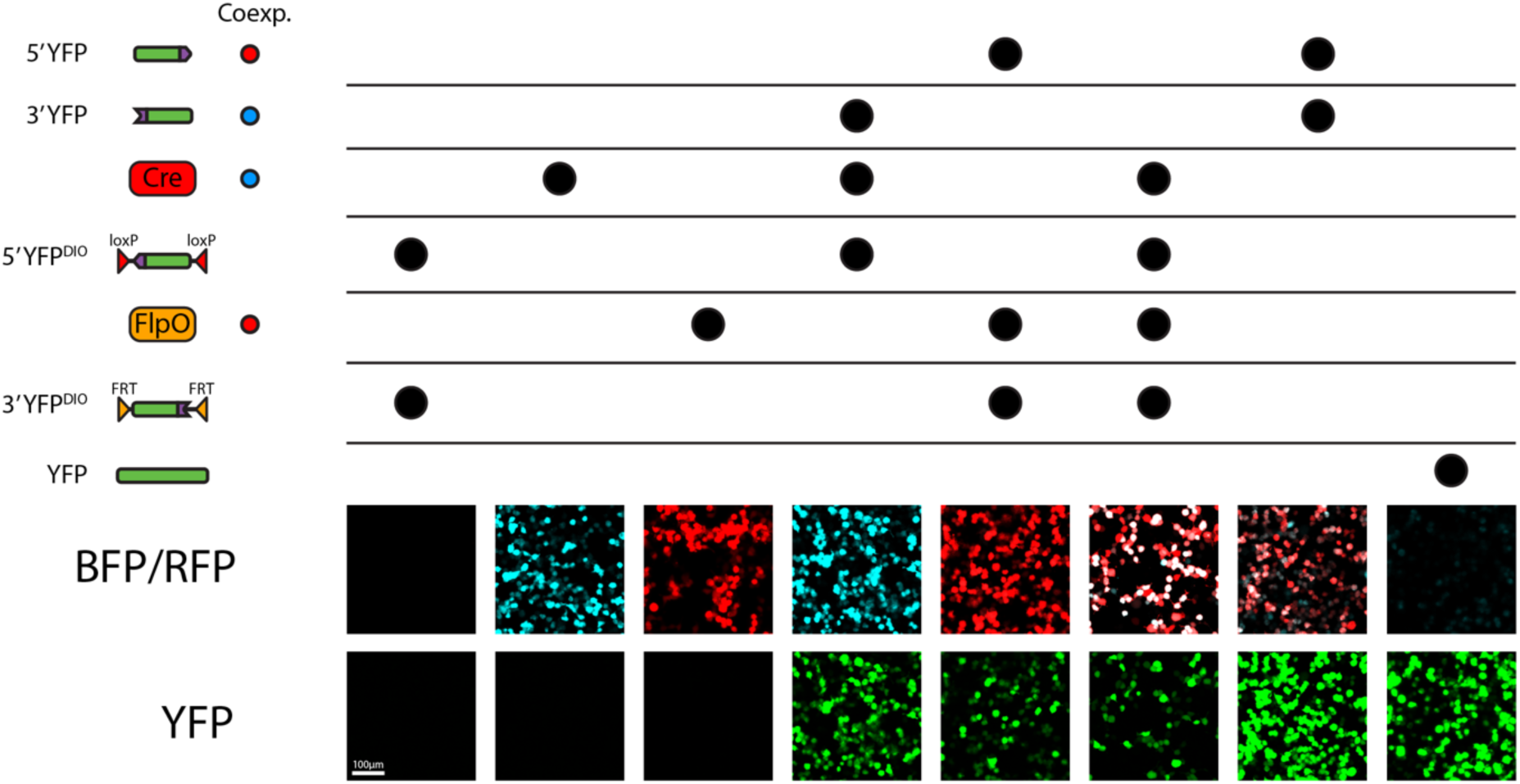
Component controls for recombinase-gated REJ-mediated YFP reconstitution. Representative fluorescence images of HEK293T cells transfected with the indicated combinations of split-YFP REJ constructs and recombinases to test Cre- and FlpO-dependent reporter activation. Columns show cells receiving 5′ YFP-DIO plus 3′ YFP-DIO; Cre alone; FlpO alone; Cre with 5′ YFP-DIO and constitutive 3′ YFP; FlpO with constitutive 5′ YFP and 3′ YFP-DIO; Cre plus FlpO with 5′ YFP-DIO and 3′ YFP-DIO; constitutive 5′ YFP plus constitutive 3′ YFP; or full-length YFP. Co-expression reporters encoded by the indicated plasmids are shown in the BFP/RFP row, and YFP reporter output is shown in the YFP row. YFP was detected when compatible 5′ and 3′ YFP RNA components were co-expressed directly, or when recombinase activity enabled expression of the corresponding DIO-gated split-YFP component. The Cre/FlpO-gated condition demonstrates combinatorial activation of YFP by recombinase-dependent expression of both REJ-compatible split-YFP RNAs.

**Figure S4.**
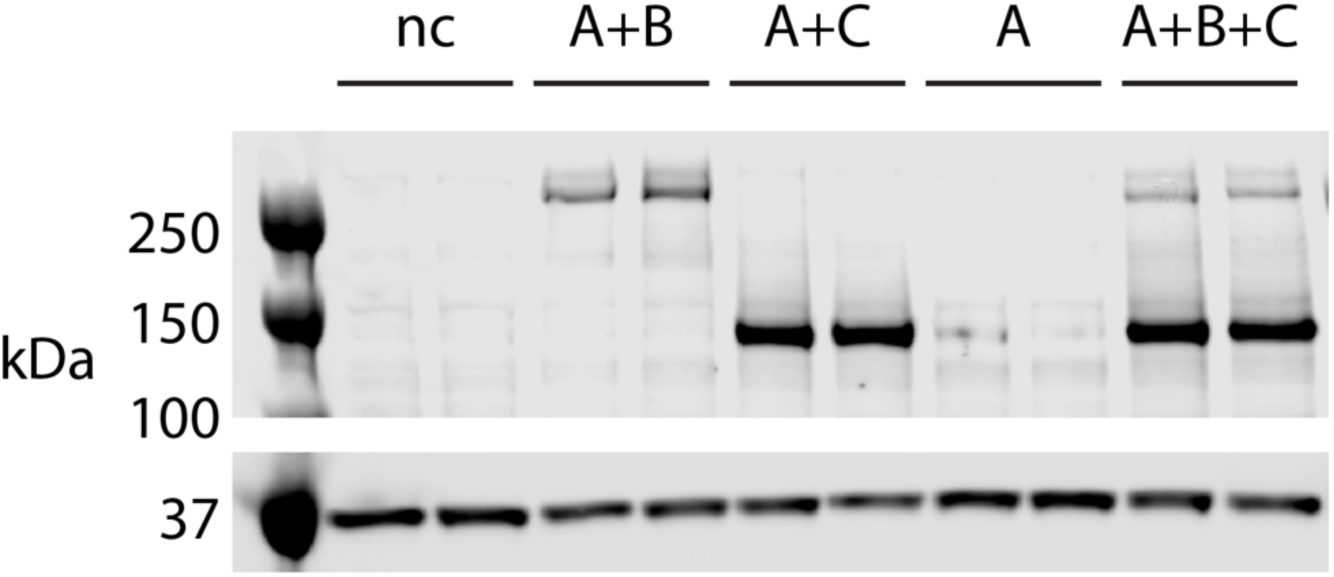
A shared 5′ REJ RNA can generate distinct protein outputs with alternative 3′ REJ partners. Western blot analysis of HEK293T cells transfected with combinations of REJ constructs designed to test whether a common 5′ RNA can splice with multiple compatible 3′ RNA partners. Construct A encodes 5′ PCDH15, construct B encodes 3′ PCDH15, and construct C encodes a YFP-containing 3′ partner. Cells were transfected with no REJ constructs, A+B, A+C, A alone, or A+B+C. Co-expression of A+B produced an approximately 260 kDa PCDH15 band, whereas co-expression of A+C produced an approximately 140 kDa PCDH15–YFP chimeric band. A alone did not produce a detectable product. Co-expression of A+B+C generated both the approximately 260 kDa PCDH15 band and the approximately 140 kDa PCDH15–YFP chimeric band, indicating that a shared 5′ REJ RNA can be routed to alternative 3′ REJ partners to produce distinct protein outputs. These results support the molecular basis for IMPLIES-style REJ logic in which one upstream RNA can enable multiple downstream outputs.

## References

1. Peterson, M.G., and Baichwal, V.R. (1993). Transcription factor based therapeutics: drugs of the future? Trends Biotechnol 11, 11–18. 10.1016/0167-7799(93)90069-l.

2. Zhang, F., Wen, Y., and Guo, X. (2014). CRISPR/Cas9 for genome editing: progress, implications and challenges. Hum Mol Genet 23, R40–46. 10.1093/hmg/ddu125.

3. Cai, M., and Yang, Y. (2014). Targeted genome editing tools for disease modeling and gene therapy. Curr Gene Ther 14, 2–9. 10.2174/156652321402140318165450.

4. Doudna, J.A. (2020). The promise and challenge of therapeutic genome editing. Nature 578, 229–236. 10.1038/s41586-020-1978-5.

5. Bains, S., Giudicessi, J.R., Odening, K.E., and Ackerman, M.J. (2024). State of Gene Therapy for Monogenic Cardiovascular Diseases. Mayo Clin Proc 99, 610–629. 10.1016/j.mayocp.2023.11.003.

6. Mai, A. (2007). The therapeutic uses of chromatin-modifying agents. Expert Opin Ther Targets 11, 835–851. 10.1517/14728222.11.6.835.

7. Cartegni, L., and Krainer, A.R. (2003). Correction of disease-associated exon skipping by synthetic exon-specific activators. Nat Struct Biol 10, 120–125. 10.1038/nsb887.

8. Will, C.L., and Lührmann, R. (2011). Spliceosome structure and function. Cold Spring Harb Perspect Biol 3. 10.1101/cshperspect.a003707.

9. Ule, J., and Blencowe, B.J. (2019). Alternative Splicing Regulatory Networks: Functions, Mechanisms, and Evolution. Mol Cell 76, 329–345. 10.1016/j.molcel.2019.09.017.

10. Wang, E.T., Sandberg, R., Luo, S., Khrebtukova, I., Zhang, L., Mayr, C., Kingsmore, S.F., Schroth, G.P., and Burge, C.B. (2008). Alternative isoform regulation in human tissue transcriptomes. Nature 456, 470–476. 10.1038/nature07509.

11. Blumenthal, T., and Thomas, J. (1988). Cis and trans mRNA splicing in C. elegans. Trends Genet 4, 305–308. 10.1016/0168-9525(88)90107-2.

12. Fiflis, D.N., Rey, N.A., Venugopal-Lavanya, H., Sewell, B., Mitchell-Dick, A., Clements, K.N., Milo, S., Benkert, A.R., Rosales, A., Fergione, S., and Asokan, A. (2024). Repurposing CRISPR-Cas13 systems for robust mRNA trans-splicing. Nat Commun 15, 2325. 10.1038/s41467-024-46172-4.

13. Puttaraju, M., Jamison, S.F., Mansfield, S.G., Garcia-Blanco, M.A., and Mitchell, L.G. (1999). Spliceosome-mediated RNA trans-splicing as a tool for gene therapy. Nat Biotechnol 17, 246–252. 10.1038/6986.

14. Sharp, P.A. (1987). Trans splicing: variation on a familiar theme? Cell 50, 147–148. 10.1016/0092-8674(87)90207-8.

15. Wally, V., Murauer, E.M., and Bauer, J.W. (2012). Spliceosome-mediated trans-splicing: the therapeutic cut and paste. J Invest Dermatol 132, 1959–1966. 10.1038/jid.2012.101.

16. Kolesnik, V.V., Nurtdinov, R.F., Oloruntimehin, E.S., Karabelsky, A.V., and Malogolovkin, A.S. (2024). Optimization strategies and advances in the research and development of AAV-based gene therapy to deliver large transgenes. Clin Transl Med 14, e1607. 10.1002/ctm2.1607.

17. Hirsch, M.L., Wolf, S.J., and Samulski, R.J. (2016). Delivering Transgenic DNA Exceeding the Carrying Capacity of AAV Vectors. Methods Mol Biol 1382, 21–39. 10.1007/978-1-4939-3271-9_2.

18. Riedmayr, L.M., Hinrichsmeyer, K.S., Thalhammer, S.B., Mittas, D.M., Karguth, N., Otify, D.Y., Böhm, S., Weber, V.J., Bartoschek, M.D., Splith, V., et al. (2023). mRNA trans-splicing dual AAV vectors for (epi)genome editing and gene therapy. Nat Commun 14, 6578. 10.1038/s41467-023-42386-0.

19. Trapani, I., Colella, P., Sommella, A., Iodice, C., Cesi, G., de Simone, S., Marrocco, E., Rossi, S., Giunti, M., Palfi, A., et al. (2014). Effective delivery of large genes to the retina by dual AAV vectors. EMBO Mol Med 6, 194–211. 10.1002/emmm.201302948.

20. Hertel, K.J., Lynch, K.W., and Maniatis, T. (1997). Common themes in the function of transcription and splicing enhancers. Curr Opin Cell Biol 9, 350–357. 10.1016/s0955-0674(97)80007-5.

21. Song, Y., Lou, H.H., Boyer, J.L., Limberis, M.P., Vandenberghe, L.H., Hackett, N.R., Leopold, P.L., Wilson, J.M., and Crystal, R.G. (2009). Functional cystic fibrosis transmembrane conductance regulator expression in cystic fibrosis airway epithelial cells by AAV6.2-mediated segmental trans-splicing. Hum Gene Ther 20, 267–281. 10.1089/hum.2008.173.

22. Pergolizzi, R.G., Ropper, A.E., Dragos, R., Reid, A.C., Nakayama, K., Tan, Y., Ehteshami, J.R., Coleman, S.H., Silver, R.B., Hackett, N.R., et al. (2003). In vivo trans-splicing of 5’ and 3’ segments of pre-mRNA directed by corresponding DNA sequences delivered by gene transfer. Mol Ther 8, 999–1008. 10.1016/j.ymthe.2003.08.022.

23. Garcia-Blanco, M.A., Puttaraju, M., Mansfield, S.G., and Mitchell, L.G. (2001). Spliceosome-mediated RNA trans-splicing in gene therapy and genomics. Gene Therapy and Regulation 1, 141–164.

24. Haddrick, M., Lear, A.L., Cann, A.J., and Heaphy, S. (1996). Evidence that a kissing loop structure facilitates genomic RNA dimerisation in HIV-1. J Mol Biol 259, 58–68. 10.1006/jmbi.1996.0301.

25. Le Hir, H., Nott, A., and Moore, M.J. (2003). How introns influence and enhance eukaryotic gene expression. Trends Biochem Sci 28, 215–220. 10.1016/s0968-0004(03)00052-5.

26. Soret, J., Gabut, M., and Tazi, J. (2006). SR proteins as potential targets for therapy. Prog Mol Subcell Biol 44, 65–87. 10.1007/978-3-540-34449-0_4.

27. Chasin, L.A. (2007). Searching for splicing motifs. Adv Exp Med Biol 623, 85–106. 10.1007/978-0-387-77374-2_6.

28. Lim, C.S., and Brown, C.M. (2016). Hepatitis B virus nuclear export elements: RNA stem-loop α and β, key parts of the HBV post-transcriptional regulatory element. RNA Biol 13, 743–747. 10.1080/15476286.2016.1166330.

29. Nott, A., Le Hir, H., and Moore, M.J. (2004). Splicing enhances translation in mammalian cells: an additional function of the exon junction complex. Genes Dev 18, 210–222. 10.1101/gad.1163204.

30. Shaul, O. (2017). How introns enhance gene expression. Int J Biochem Cell Biol 91, 145–155. 10.1016/j.biocel.2017.06.016.

31. Patel, A., Zhao, J., Duan, D., and Lai, Y. (2019). Design of AAV Vectors for Delivery of Large or Multiple Transgenes. Methods Mol Biol 1950, 19–33. 10.1007/978-1-4939-9139-6_2.

32. Chamberlain, K., Riyad, J.M., and Weber, T. (2016). Expressing Transgenes That Exceed the Packaging Capacity of Adeno-Associated Virus Capsids. Hum Gene Ther Methods 27, 1–12. 10.1089/hgtb.2015.140.

33. Zaydon, Y.A., and Tsang, S.H. (2024). The ABCs of Stargardt disease: the latest advances in precision medicine. Cell Biosci 14, 98. 10.1186/s13578-024-01272-y.

34. Li, R., Jing, Q., She, K., Wang, Q., Jin, X., Zhao, Q., Su, J., Song, L., Fu, J., Wu, X., et al. (2023). Split AAV8 Mediated ABCA4 Expression for Gene Therapy of Mouse Stargardt Disease (STGD1). Hum Gene Ther 34, 616–628. 10.1089/hum.2023.017.

35. Trapani, I. (2018). Dual AAV Vectors for Stargardt Disease. Methods Mol Biol 1715, 153–175. 10.1007/978-1-4939-7522-8_11.

36. Lindley, S.R., Subbaiah, K.C.V., Priyanka, F., Poosala, P., Ma, Y., Jalinous, L., West, J.A., Richardson, W.A., Thomas, T.N., and Anderson, D.M. (2024). Ribozyme-activated mRNA trans-ligation enables large gene delivery to treat muscular dystrophies. Science 386, 762–767. 10.1126/science.adp8179.

37. Lin, J., Lin, Y., Liu, N., Cao, W., Zhang, J., Wen, S., Zhang, Y., Liao, W., Hong, Z., Lin, Y., et al. (2026). AAVLINK: A potent DNA-recombination method for large cargo delivery in gene therapy. Cell 189, 969–986.e917. 10.1016/j.cell.2025.12.039.

38. Hur, S. (2019). Double-Stranded RNA Sensors and Modulators in Innate Immunity. Annu Rev Immunol 37, 349–375. 10.1146/annurev-immunol-042718-041356.

39. Akira, S., Uematsu, S., and Takeuchi, O. (2006). Pathogen recognition and innate immunity. Cell 124, 783–801. 10.1016/j.cell.2006.02.015.

40. Li, D., and Wu, M. (2021). Pattern recognition receptors in health and diseases. Signal Transduction and Targeted Therapy 6, 291. 10.1038/s41392-021-00687-0.

41. Haery, L., Deverman, B.E., Matho, K.S., Cetin, A., Woodard, K., Cepko, C., Guerin, K.I., Rego, M.A., Ersing, I., Bachle, S.M., et al. (2019). Adeno-Associated Virus Technologies and Methods for Targeted Neuronal Manipulation. Front Neuroanat 13, 93. 10.3389/fnana.2019.00093.

42. Khan, M.S., Khalid, A.M., and Malik, K.A. (2005). Intein-mediated protein trans-splicing and transgene containment in plastids. Trends Biotechnol 23, 217–220. 10.1016/j.tibtech.2005.03.006.

43. Tornabene, P., Trapani, I., Minopoli, R., Centrulo, M., Lupo, M., de Simone, S., Tiberi, P., Dell’Aquila, F., Marrocco, E., Iodice, C., et al. (2019). Intein-mediated protein trans-splicing expands adeno-associated virus transfer capacity in the retina. Sci Transl Med 11. 10.1126/scitranslmed.aav4523.

44. Tasfaout, H., Halbert, C.L., McMillen, T.S., Allen, J.M., Reyes, T.R., Flint, G.V., Grimm, D., Hauschka, S.D., Regnier, M., and Chamberlain, J.S. (2024). Split intein-mediated protein trans-splicing to express large dystrophins. Nature 632, 192–200. 10.1038/s41586-024-07710-8.

45. Duan, D., Yue, Y., and Engelhardt, J.F. (2001). Expanding AAV packaging capacity with trans-splicing or overlapping vectors: a quantitative comparison. Mol Ther 4, 383–391. 10.1006/mthe.2001.0456.

46. Veitia, R.A. (2007). Exploring the molecular etiology of dominant-negative mutations. Plant Cell 19, 3843–3851. 10.1105/tpc.107.055053.

47. Duan, D. (2023). Lethal immunotoxicity in high-dose systemic AAV therapy. Mol Ther 31, 3123–3126. 10.1016/j.ymthe.2023.10.015.

48. Cottrell, K.A., Andrews, R.J., and Bass, B.L. (2024). The competitive landscape of the dsRNA world. Mol Cell 84, 107–119. 10.1016/j.molcel.2023.11.033.

49. Chan, C.P., and Jin, D.Y. (2022). Cytoplasmic RNA sensors and their interplay with RNA-binding partners in innate antiviral response: theme and variations. Rna 28, 449–477. 10.1261/rna.079016.121.

50. Im, J.H., Duic, I., Yoshimura, S.H., Onomoto, K., Yoneyama, M., Kato, H., and Fujita, T. (2023). Mechanisms of length-dependent recognition of viral double-stranded RNA by RIG-I. Sci Rep 13, 6318. 10.1038/s41598-023-33208-w.

51. Liu, G., and Gack, M.U. (2020). Distinct and Orchestrated Functions of RNA Sensors in Innate Immunity. Immunity 53, 26–42. 10.1016/j.immuni.2020.03.017.

52. del Toro Duany, Y., Wu, B., and Hur, S. (2015). MDA5-filament, dynamics and disease. Curr Opin Virol 12, 20–25. 10.1016/j.coviro.2015.01.011.

53. McGarry, N., Murray, C.L., Garvey, S., Wilkinson, A., Tortorelli, L., Ryan, L., Hayden, L., Healy, D., Griffin, E.W., Hennessy, E., et al. (2021). Double stranded RNA drives innate immune responses, sickness behavior and cognitive impairment dependent on dsRNA length, IFNAR1 expression and age. bioRxiv. 10.1101/2021.01.09.426034.

54. Schweinoch, D., Bachmann, P., Clausznitzer, D., Binder, M., and Kaderali, L. (2020). Mechanistic modeling explains the dsRNA length-dependent activation of the RIG-I mediated immune response. J Theor Biol 500, 110336. 10.1016/j.jtbi.2020.110336.

55. McClements, M.E., and MacLaren, R.E. (2017). Adeno-associated Virus (AAV) Dual Vector Strategies for Gene Therapy Encoding Large Transgenes. Yale J Biol Med 90, 611–623.

56. Li, J., Sun, W., Wang, B., Xiao, X., and Liu, X.Q. (2008). Protein trans-splicing as a means for viral vector-mediated in vivo gene therapy. Hum Gene Ther 19, 958–964. 10.1089/hum.2008.009.

57. Loonstra, A., Vooijs, M., Beverloo, H.B., Allak, B.A., van Drunen, E., Kanaar, R., Berns, A., and Jonkers, J. (2001). Growth inhibition and DNA damage induced by Cre recombinase in mammalian cells. Proc Natl Acad Sci U S A 98, 9209–9214. 10.1073/pnas.161269798.

58. Trapani, I. (2019). Adeno-Associated Viral Vectors as a Tool for Large Gene Delivery to the Retina. Genes (Basel) 10. 10.3390/genes10040287.

59. Cullis, P.R., and Hope, M.J. (2017). Lipid Nanoparticle Systems for Enabling Gene Therapies. Mol Ther 25, 1467–1475. 10.1016/j.ymthe.2017.03.013.

60. Zabin, I., and Villarejo, M.R. (1975). Protein complementation. Annu Rev Biochem 44, 295–313. 10.1146/annurev.bi.44.070175.001455.

